# Co-option of immune effectors by the hormonal signalling system triggering metamorphosis in *Drosophila melanogaster*

**DOI:** 10.1101/2021.05.22.444626

**Authors:** Catarina Nunes, Takashi Koyama, Élio Sucena

**Affiliations:** Evolution and Development Laboratory, Instituto Gulbenkian de Ciência, Oeiras, Portugal; Section for Cell and Neurobiology, Department of Biology University of Copenhagen, Copenhagen, Denmark; Departamento de Biologia Animal, Faculdade de Ciências, Universidade de Lisboa, Lisbon, Portugal

**Author notes:** Corresponding authors (CN); (ES).

**Keywords:** Ecdysone, Anti-Microbial Peptides (AMP), Insect Metamorphosis, Drosomycin

## Abstract

Insect metamorphosis is regulated by the production, secretion and degradation of two peripheral hormones: 20-hydroxyecdysone (ecdysone) and juvenile hormone (JH). In addition to their roles in developmental regulation, increasing evidence suggests that these hormones are involved in innate immunity processes, such as phagocytosis and the induction of antimicrobial peptide (AMP) production. AMP regulation includes systemic responses as well as local responses at surface epithelia that contact with the external environment. At pupariation, *Drosophila melanogaster* increases dramatically the expression of three AMP genes, *drosomycin (drs), drosomycin-like 2 (drsl2)* and *drosomycin-like 5 (drsl5)*. We show that the systemic action of *drs* at pupariation is dependent on ecdysone signalling in the fat body and operates via the ecdysone downstream target, Broad-Z4. In parallel, ecdysone also regulates local responses, specifically through the activation of *drsl2* expression in the gut. Finally, we confirm the relevance of this ecdysone dependent AMP expression for the control of bacterial persistence by showing that flies lacking *drs* expression in the fat body have higher bacterial persistence over metamorphosis. Together, our data emphasize the importance of the association between ecdysone signalling and immunity using *in vivo* studies and establish a new role for ecdysone at pupariation, which impacts developmental success by regulating the immune system in a stage-dependent manner. We speculate that this co-option of immune effectors by the hormonal system may constitute a pre-emptive mechanism to control bacterial numbers in the pupa at the core of metamorphosis evolution.

## Introduction

Insect development and physiology are mainly regulated by two peripheral hormones: the steroid hormone, 20-hydroxyecdsyone (hereafter, ecdysone) and the sesquiterpenoid hormone, juvenile hormone (JH) (1,2). In vertebrates, steroid hormones and their nuclear receptors are key regulators of systemic immune responses, namely through the enhancement of inflammation (3). Similarly, in insects, steroid hormones are also known to be involved in innate immunity (4–9). Ecdysone titre is tightly regulated to control developmental transitions and as such, steroid hormone-dependent activation of innate immunity seems appropriate for stage-specific regulation of immunity.

When in contact with a microorganism, insects rely on the activation of three tightly related immunity processes: i) cellular immunity (i.e. phagocytosis and encapsulation of invading microorganisms); ii) the induction of proteolytic cascades, that culminate in melanisation; and iii) production of antimicrobial peptides (AMPs), which is primarily regulated at the transcriptional level through two distinct signalling pathways, Toll and Imd, triggered by sensing of Lys-type peptidoglycan and β-1,3-glucan or meso-diaminopimelic acid (DAP)-type peptidoglycan, respectively (10–12). In insects with complete metamorphosis (holometabolans), AMP genes are usually silent in the absence of an immune challenge and their expression is induced upon injury and/or infection (13). However, indirect evidence suggests that AMPs may be up-regulated at specific developmental stages irrespective of pathogen presence. Studies in the moth *Manduca sexta* show that expression of lysozyme and of the AMPs *cecropin A, cecropin B* and *hemolin* increases dramatically during metamorphosis, although it is not clear what regulates this increase and its role(s) *in vivo* (14–16). This transcriptional AMP peak occurs before the end of histolysis of the larval midgut, that is, before cells are potentially exposed to any pathogen or bacterium. In this context, the accumulation of AMPs at the end of the feeding stage may constitute a first response against the possible harmful effects to the pupa of expanding bacteria carried-over from the gut remodelling process (17–20). Nonetheless, studies in dipterans show that some species of bacteria in the larval gut can persist inside the host throughout metamorphosis (19–22). In *Drosophila melanogaster*, RNAseq analysis revealed a peak in the expression of AMPs at pupariation, of which around 95% consists of three AMPs: *drs, drosomycin-like 2 (drsl2)* and *drosomycin-like 5 (drsl5)* (23). Furthermore, five different AMPs (*diptericin, drs, attacin-A, metchnikowin* and *cecropin A1*) are shown to contain putative *cis*-regulatory elements for potential binding of the functional ecdysone receptor complex, suggesting an ecdysone-dependent control of their expression (24). Importantly, the compelling association between ecdysone signalling and immunity at this particular stage of development is not restricted to the humoral arm of the immune response. An ecdysone-dependent increase in phagocytic capacity of plasmatocytes at pupariation has been described and hypothesized to be particularly important given the higher predisposition for infection at the pupal stage (6).

Considering the above, we hypothesized that ecdysone signalling controlling development was co-opted to regulate the expression of AMPs at pupariation. Previous *in vitro* study has shown that ecdysone promotes humoral immunity by increasing the expression of AMP genes, such as *diptericin, cecropin* and *attacin*, via the EcR-USP receptor complex in S2 cells (5). In addition, a microarray analysis of 20E-treated S2 cells revealed the existence of two distinct mechanisms for 20E-regulated AMP expression: one in which the critical step is the activation of peptidoglycan recognition protein (PGRP)-LC - a critical pattern recognition receptor (PRR) of the Imd pathway - by ecdysone for controlling the expression of *cecropin A1, attacin-A and defensin*; and another, in which the hormonally-controlled mechanism is absolutely required for the expression of *drs, metchnikowin* and *diptericin* (4). However, the importance of the requirement of ecdysone for the expression of AMPs, and its role, nor the regulatory mechanism through which this hormone affects innate immunity have been explored.

In this study, we demonstrate that ecdysone regulates the expression of AMPs *in vivo* at pupariation, both systemically (by regulating the expression of *drs*) and locally in the midgut (by regulating the expression of *drsl2*). Importantly, the systemic expression of *drs* has an impact on the number of bacteria that persists and proliferates during metamorphosis, corroborating the hypothesis that the AMP peak has a function in controlling bacteria during metamorphosis. Furthermore, we show that this association between the expression of *drs* and ecdysone in the absence of infection occurs solely at pupariation and through the early ecdysone-response gene Broad-Z4, which is only expressed at the end of the feeding stage (25). Taken together, our findings fill a gap in current knowledge by shedding light onto the mechanisms and biological significance of hormonal control of humoral immunity at metamorphosis with potential evolutionary importance.

## Results

### *drs, drsl2* and *drsl5* are expressed at pupariation in a bacteria-independent manner

Around 95% of the anti-microbial peptide (AMP) peak at pupariation, mentioned above (23), consists of three AMPs: *drs, drsl2* and *drsl5* (Supplementary Figure 1). Upon gut remodelling at metamorphosis, an ineffective gut purge could lead to the canonical activation of immunity pathways, resulting in a peak of AMP expression. To test if the presence of bacteria constitutes the trigger for the expression of *drs, drsl2* and *drsl5*, we performed real-time quantitative PCR (qPCR) in mid-L3 larvae (L3) and newly pupariated (P0) samples of germ-free (GF) or standardly-raised individuals. We confirmed that expression increases at pupariation for all AMPs tested (Figure 1, S1 table, Supplementary Figure 2), including *drs (lsmean, pairwise∼{w-};+;+* | *Stage*drs<0*.*0001), drsl2 (lsmeans(pairwise∼{w};+;+*|*Stage*drsl2 <0*.*0001))* and *drsl5 (lsmeans(pairwise∼{w-};+;+*|*Stage*drsl5<0*.*000))*. This increment is not lost under GF conditions with the putative exceptions of *cecropinA1* and *defensin*. Importantly, the three main AMPs under scrutiny, *drs, drsl2* and *drsl5*, show a marked independence of the presence of microbes, including yeast, for their expression at this developmental transition (*lsmeans(pairwise∼{w};+;+*|*Stage*drs <0*.*0001); lsmeans(pairwise∼{w-}; +;+*|*Stage*drsl2<0*.*0001); lsmeans(pairwise∼{w};+;+*|*Stage*drsl5 <0*.*0001))* (Figure 1, S1 table). In turn, this is not the case for other AMPs, such as for example *cecropinA1* or *defensin*, which show decreased expression in the absence of gut bacteria (Supplementary Figure 2, S1 table).

**Figure 1.**
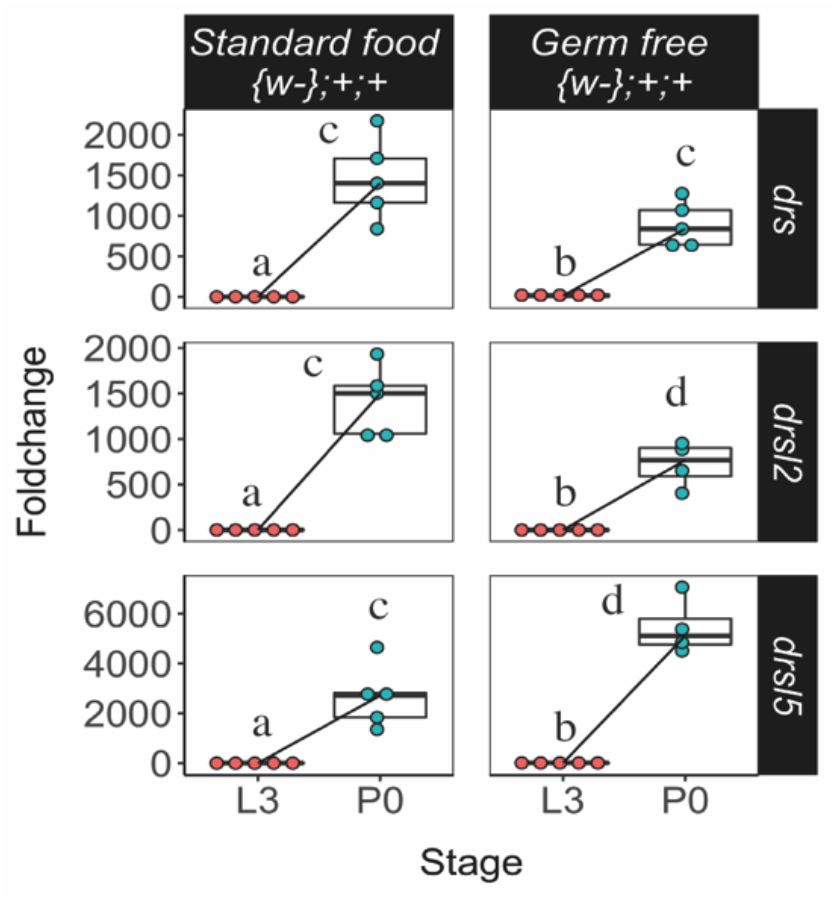
Expression of *drs, drsl2* and *drsl5* is independent of bacterial presence. The expression levels of *drs, drsl2* and *drsl5* increase at pupariation and are not lost under GF conditions. Different letters represent statistically significant differences in foldchange. Foldchange was determined using the ΔΔCT method.

Together, these results reject the hypothesis that expression of the three main AMPs at pupariation is dependent on microbial induction and suggest that it may be hardwired to the developmental programme.

### Ecdysone regulates the expression of *drs*

We focused primarily on two main sources of AMP expression with systemic action, the fat body and haemocytes (12). We manipulated ecdysone signalling sensitivity in these tissues, simultaneously and specifically, to ensure no perturbation of normal development and pupariation timing. For that we used *Cg-Gal4* combined with a *tub-Gal80[ts]*, a thermo-sensitive allele of *Gal80* under the control of the tubulin promoter (hereafter *Cg[ts]*), to drive the expression of a dominant negative ecdysone receptor under the control of the UAS sequence (26) *(EcR-A DN W650A TP3*, hereafter *EcRDN*). The reduction of ecdysone sensitivity was triggered from the onset of L3 or at wandering stage, and resulted in the elimination of the *drs* peak at P0, compared to control conditions (Figure 2A, (*lsmeans(pairwise∼Stage*|*Cg[ts]>mcD8GFP)=0*.*0001*, lsmeans(pairwise∼Stage|*Cg[ts] >EcRDN) =*0.0017, lsmeans(pairwise∼Genotype|P0) <0.0001)). However, the expression of *drsl2* and *drsl5* (Figure 2A, S2 Table) and of the other AMPs (Supplementary Figure 3-1, S2 Table) was not eliminated. Furthermore, *drs* expression still increases in mutant flies for *dif*, the NF-κB factor that triggers *drs* expression upon activation of the Toll pathway (Supplementary Figure 3-2, lsmeans(pairwise∼Stage|*{w-};+;+<*0.0001, *lsmeans(pairwise∼Stage*|*{w-};dif1;+*)<0.0001, *lsmeans(pairwise∼Genotype*|*P0)= 0*.*0703*)), suggesting that the expression of *drs* at pupariation is partially independent of the action of the NF-κB factor.

**Figure 2.**
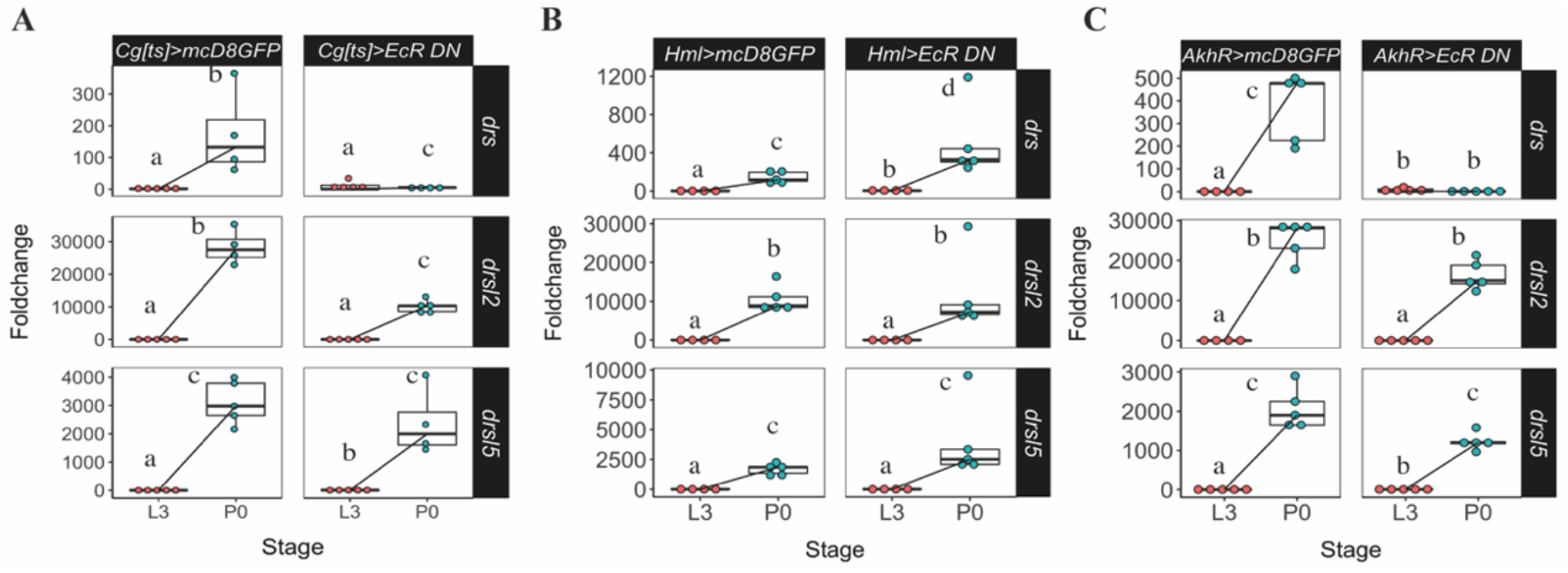
Expression of drs at pupariation is developmentally regulated by ecdysone in the fat body. (A) Foldchange increase in drs expression (but not of *drsl2* or *drsl5*) at pupariation was eliminated when ecdysone sensitivity is decreased specifically in the fat body and haemocytes simultaneously. (B) Reduced ecdysone sensitivity in the haemocytes does not affect the expression of the AMPs under study. (C) Reduced ecdysone signalling in the fat body resulted in decreased *drs* expression at pupariation (lsmeans(pairwise∼Stage|AkhR>mcD8GFP <0.0001, lsmeans(pairwise∼Stage|AkhR>EcRDN)=0.6709), without affecting the expression of *drsl2* and *drsl5* (lsmeans (pairwise∼Stage|AkhR>mcD8GFP)<0.0001, lsmeans(pairwise∼Stage|AkhR>EcRDN) <0.0001), for both genes). Different letters represent statistically significant differences in foldchange. Foldchange was determined using the ΔΔCT method.

This result was further confirmed by targeting haemocytes and fat body independently, using the specific drivers *Hemolectin-Gal4* (*Hml-ΔGal4*) for the haemocytes and *Adipokinetic hormone receptor-Gal4* (*AkhR-Gal4)* for the fat body. When ecdysone sensitivity was reduced in the haemocytes, expression of *drs, drsl2* and *drsl5*, as well as of other four AMPs, was unaffected (*lsmeans(pairwise∼Stage*|*Hml>mcD8GFP)<0*.*0001, lsmeans(pairwise∼Stage*|*Hml>EcR DN)<0*.*0001*), for the three genes), (Figure 2B and Supplementary Figure 3-3, S3 Table). However, although reduced ecdysone sensitivity in the fat body (*AkhR>EcRDN*) did not affect the expression of *drsl2* and *drsl5* (Figure 2C, (*lsmeans(pairwise∼Stage*|*AkhR >mcD8GFP) <0*.*0001, lsmeans (pairwise∼Stage*|*AkhR>EcRDN)<0*.*0001)*, for both genes) or of other AMPs (Supplementary Figure 3-4, S4 Table), it resulted in decreased expression of *drs* at pupariation. (Figure 2C, *lsmeans(pairwise∼Stage*|*AkhR>mcD8GFP) <0*.*0001, lsmeans(pairwise∼Stage*|*AkhR >EcRDN) =0*.*6709*). It is noteworthy to mention that this reduction of *drs* expression is not likely due to the EcR-mediated disruption of fat body metabolism or abnormal developmental effects because most of the AMP gene expression was not altered.

Altogether, these results indicate that *drs* expression - but not *drsl2* or *drsl5 -* is systemically regulated by ecdysone signalling specifically in the fat body at pupariation.

### Expression of *drsl2* in the midgut is regulated by Ecdysone

Having determined that the expression of *drsl2* and *drsl5* is developmentally regulated but does not involve the fat body, we turned our attention to two of the most important tissues for local immune response: the epithelial layers of the gut and trachea (27). First, we overexpressed *EcRDN* specifically in the most abundant cells of the midgut, the enterocytes, using the specific driver *Mex-Gal4* (*Mex*>*EcRDN*). Unlike the fat body, the gut is not responding to the pupariation ecdysone pulse by increasing *drs* expression (Figure 3A). In contrast, decreased ecdysone sensitivity specifically in the midgut resulted in a drastic reduction (*circa* 600-fold) of *drsl2* expression, very close to the L3 basal levels (Figure 3A), *lsmeans(pairwise∼Genotype*|*P0)<0*.*0001; lsmeans(pairwise∼Stage*|*Mex> EcRDN) <0*.*0001*)).

**Figure 3.**
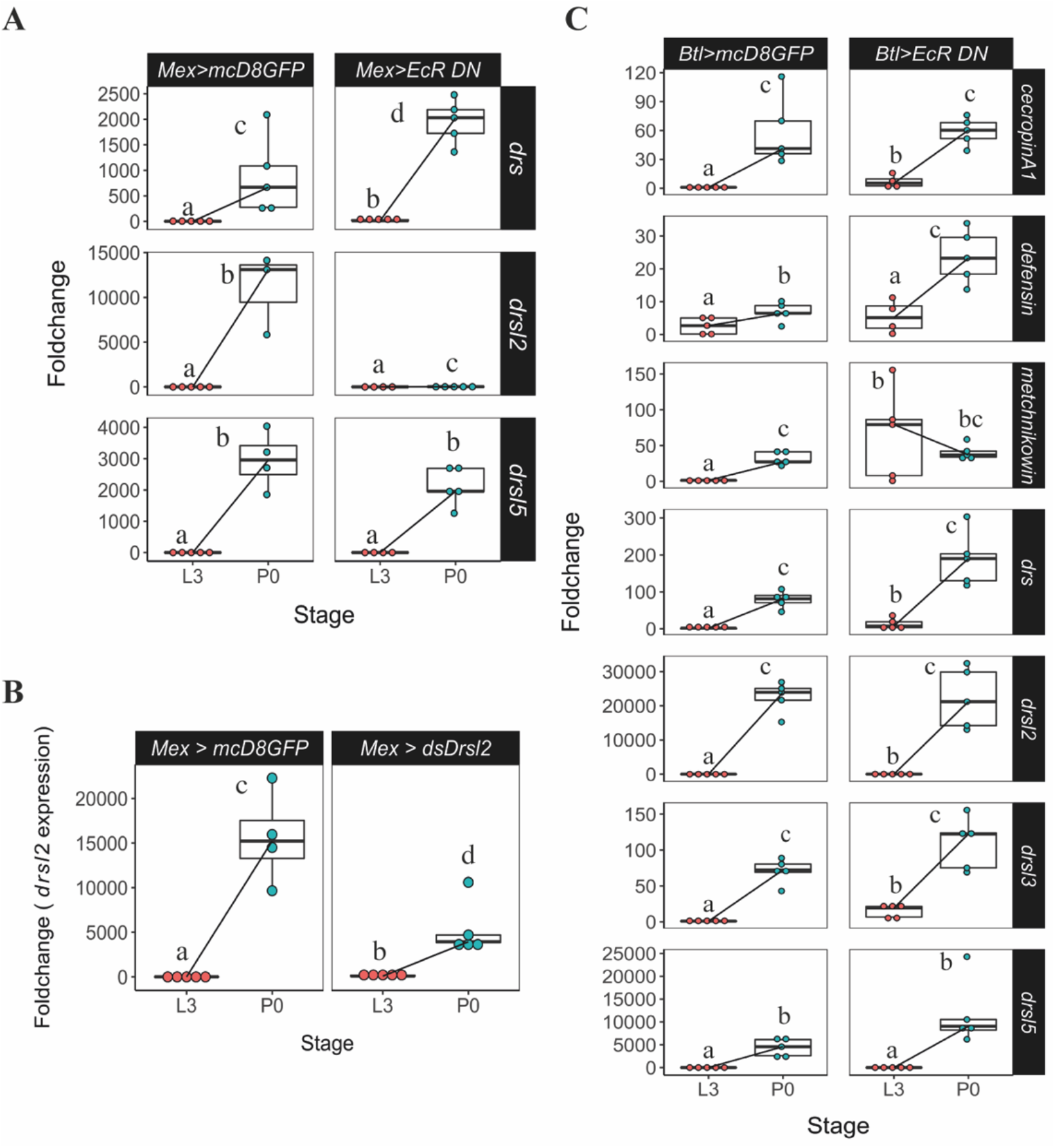
Gut and tracheal expression of relevant AMPs. (A) Manipulation of ecdysone sensitivity specifically in the midgut reduced the expression of *drsl2*, but not of *drs* or *drsl5*. (B) Knock-down of *drsl2* specifically in the midgut resulted in an overall decrease in the expression of this AMP. Each dot represents a sample of five pooled individuals; the lines connect the median of the samples at L3 and P0. (C) Ecdysone signalling in the trachea does not affect the expression of any AMP under study. Different letters represent statistically significant differences in foldchange. Foldchange was determined using the ΔΔCT method.

We further confirmed this result by inducing RNAi (through the expression of double stranded RNA for *drsl2* under UAS control) and consequent knock-down of *drsl2* specifically in the midgut (*Mex>ds-drsl2*). This resulted in an overall decrease in the expression of this AMP at pupariation (Figure 3B, *lsmeans(pairwise∼Genotype*|*P0)<0*.*0237; lsmeans(pairwise∼Stage*|*Mex>ds-drsl2) <0*.*0001*), which indicates that the midgut is a source of the ecdysone-dependent production of *drsl2* at this stage. Interestingly, we failed again to detect any difference in the expression of *drsl5*, one of the three main players in the AMP ecdysone-dependent peak (Figure 3A, *lsmeans (pairwise∼Genotype*|*P0)<0*.*4479; lsmeans(pairwise∼Stage*|*Mex>EcRDN)<0*.*0001*), S5 Table). However, other tested AMPs also appear to be, at least partially, regulated by ecdysone signalling in the midgut at pupariation.

Under ecdysone-signalling down-regulation conditions, significant decrease in mRNA levels were observed for: *cecropin A1* (*lsmeans(pairwise∼Genotype*|*P0) <0*.*0001; lsmeans(pairwise∼Stage*|*Mex>EcRDN)=0*.*0038), defensin (lsmeans (pairwise∼Genotype*|*P0)<0*.*0001; lsmeans(pairwise∼Stage*|*Mex>EcR DN)=0*.*3866*),*metchnikowin* (*lsmeans(pairwise∼Genotype*|*P0)=0*.*0060; lsmeans (pairwise∼Stage*|*Mex>EcRDN<0*.*0001*), and *drsl3((lsmeans(pairwise∼Genotype*|*P0) =0*.*0008;lsmeans(pairwise∼Stage*|*Mex>EcRDN)=0*.*2084*), Supplementary Figure 4).

On the other hand, decreased ecdysone sensitivity in the trachea, using the specific driver *btl-*Gal4 (*btl>EcRDN*), did not have an impact on the expression of any of the three major AMPs under study (Figure 3C, S6 Table). These results show that a layer of local AMP action is at play through the sensing of the ecdysone peak at the onset of pupariation in the gut epithelium.

Together the local and systemic mechanisms of action that share ecdysone control, consisting of *drsl2* and *drs*, account for about 70% of the pupariation AMP peak. Contrastingly, the data also point towards an alternative mechanism for regulating the expression of *drsl5* at pupariation, which is either independent of ecdysone or is regulated by this hormone in an alternative tissue or cell type that were not tested.

### Br is involved in systemic *drs* induction through the fat body at pupariation

Next, we focused our efforts in trying to uncover more of the molecular regulation of *drs* expression, the systemic arm of this ecdysone co-opted immune action.

Previous RNAseq data showed that the expression of *drs* markedly increases at pupariation and gradually drops the expression in next 24 hours (Supplementary Figure 1, (23)). We further confirmed that the expression of *dhr3* (a gene directly regulated by the ecdysone-ecdysone receptor complex) and that of *drs* are tightly correlated and peak exclusively at pupariation (Figure 4A). Importantly, this robust expression of *drs* at the onset of metamorphosis seems to be a general feature regardless of their genetic background (Supplementary Figure 5-1).

**Figure 4.**
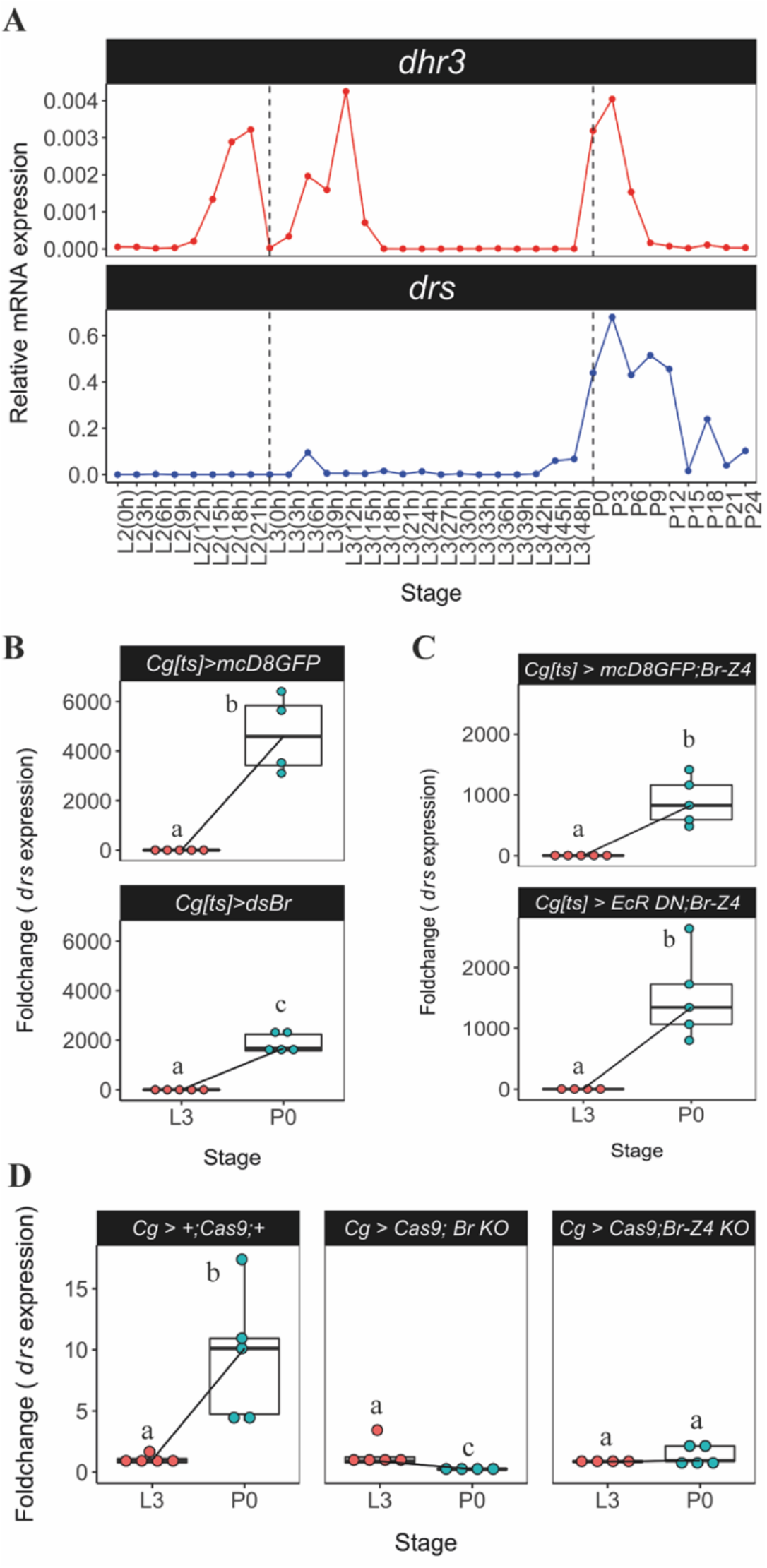
Ecdysone signalling regulates fat body expression of *drs* at pupariation via Br-Z4. **(A)** *drs* expression is temporally correlated with a proxy of ecdysone activity, *dhr3*. Samples of *D. melanogaster* larvae and pupae were precisely staged at the onset of the instars and collected every 3h. The expression of *dhr3* and *drs* was determined by qPCR for each timepoint. Dashed black lines represent the moult to the L3 or pupariation. **(B)** Knockdown of *br* specifically in the fat body and haemocytes leads to decreased expression of *drs* at P0. (**C**) Co-overexpression of *br*-*Z4* when ecdysone sensitivity is decreased in the fat body and haemocytes is sufficient to rescue the expression of *drs* at pupariation (**D**) Conditional CRISRP-mediated knock-outs for the common region of *br* and *br-Z4* lead to loss of *drs* expression at pupariation. Each dot represents a sample of five pooled individuals; the lines connect the median of the samples at L3 and P0; different letters represent statistically significant differences in foldchange, determined using the ΔΔCT method.

*br* is a direct target of ecdysone receptor, characterized as a holometabolan pupal specifier with increased expression before pupariation (25,28,29). Because the transcriptional peak of *drs* is only found at the moult from the L3 to pupa but not from the L2 to L3, Br seems a good candidate for regulation of the *drs* expression. We thus examined the association between *br* and the ecdysone-sensitive systemic induction of *drs* at metamorphosis by knocking-down *br* specifically in the fat body and haemocytes. This led to a significant decrease in the expression of *drs* at pupariation, when compared to the control (Figure 4B, *lsmeans(pairwise∼Genotype*|*P0)= 0*.*0510*)), suggesting a role for *br* in the regulation of *drs* expression. To further explore this possibility, we simultaneously reduced ecdysone signalling and restored *br* in the haemocytes and fat body. This was sufficient to rescue the expression of *drs* to control levels at P0 (Figure 4C, *lsmeans(pairwise∼Genotype*|*P0)= 0*.*3068*). Furthermore, we generated two different conditional CRISPR-mediated knock-out constructs (30,31), one targeting a common region to all *br* isoforms and, the other, a region specific to *br-Z4*, previously associated with AMP expression using *in vitro* techniques in a different species (32). Knock-out of all *br* isoforms or of *br-Z4* alone in haemocytes and fat body, led to the elimination of the *drs* peak at pupariation (Figure 4D, *lsmeans(pairwise∼Stage*|*Cg> +;Cas9;+) <0*.*0001;lsmeans(pairwise∼Stage*| *Cg>Cas9;br KO)=0*.*0001, lsmeans(pairwise∼Stage*|*Cg>Cas9;br-Z4 KO)=0*.*4691*, S8 Table), supporting that Br is involved in the regulation of *drs* expression at pupariation downstream of the ecdysone receptor complex in the fat body.

To further explore this regulation of *drs* expression by Br-Z4, we used the JASPAR (http://jaspar.genereg.net) matrix MA0013.1 to run a comprehensive search for Br-Z4 binding sites over 2 kb of putative promoter region that is upstream of the coding sequence of *drs*, obtained from Flybase (http://flybase.net/), using FIMO software (33) (Supplementary Figure 5-2). We identified three putative Br-Z4 binding sites (p-value<0.0001) in the predicted 5’ promoter region of *drs* (Supplementary Figure 5-3).

Together, these results support the idea that Br, specifically isoform Br-Z4, is involved in the regulation of the expression of *drs* either directly or indirectly at pupariation.

### *drs* is necessary to reduce bacteria derived from imperfect larval gut purge

Having establish the basis of the regulation of *drs* at the onset of metamorphosis, we set to determine its relevance to the control of bacterial numbers throughout pupal development. Indeed, before pupariation and metamorphosis, larvae purge their gut contents and any imperfection in this process creates the conditions in the pupa for a putatively rampant infection. Thus, we examined whether or not gut purge eliminates all bacteria from the gut and how this process may be dependent on the systemic action of Drs.

Under our standard rearing conditions, the mortality between P0 and pharate adult is negligible, which allows us to infer and correlate the bacterial numbers detected in each of these developmental stages (Supplementary Figure 6, S7 Table). After gut purge, approximately 5% of control P0 individuals contained bacteria in the order of the thousands and another 12% in the order of the hundreds (Figure 5A). However, at the pharate adult stage, the number of individuals in these two categories decreased alongside an increase in the number of individuals without detectable bacteria (Figure 5B). In contrast, *drs* null mutants showed an increase rather than a decrease in the number of pupae with more bacteria between these two stages (Figure 5C and 5D), supporting that Drs is necessary to eliminate gut derived bacteria during metamorphosis.

**Figure 5.**
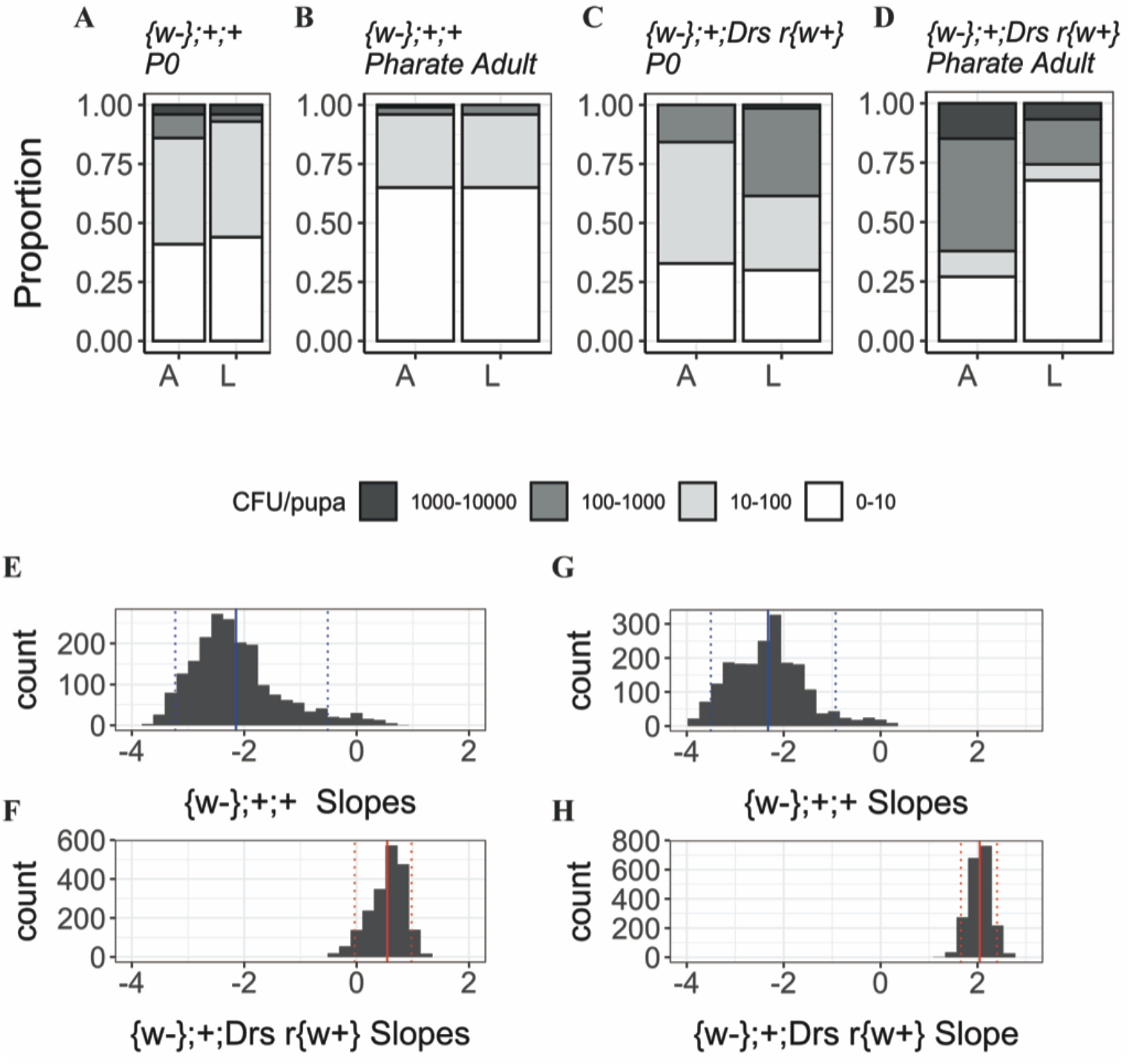
(A-F) Bacteria quantification (*Lactobacillus* (L) and *Acetobacter* (A)) in P0 pupae and pharate adults in wild-type and drs mutants. (A-B) In control flies *({w-};+;+*), the number of pupae with more bacteria decreases between P0 and pharate adult stages. (C-D) In the *drs* mutant line (*{w-};+; Drs r{w+})*, there are more individuals with higher loads of bacteria persisting from P0 to pharate adult. (E-F) Slopes for the number of *Lactobacillus* in control and the mutant lines. (E) Control samples have negative slopes indicating a decrease in the number of bacteria between stages. (F) *drs* mutant samples have positive slope values, representative of an increase in the number of bacteria across metamorphosis. (G-H) Slopes for the number of *Acetobacter* in the control and the mutant lines. (G) Control samples distribution show negative slopes, i.e., a decrease in the number of bacteria between stages. (H) *drs* null mutant displays positive slope values. Doted lines represent the 0.05 and 0.95 quantiles; full line represents the mean slope.

Using a resampling strategy, we determined that most coefficients associated with the control line were negative (Figure 5E (*Median: -2*.*279 (IQR: -2*.*682 -1*.*803); One-way Wilcoxon test for median<0: <2*.*2e-16, r= 0*.*864)* and Figure 5G *(Median: -2*.*317 (IQR: - 2*.*907 -1*.*858); One-way Wilcoxon test for median<0: <2*.*2e-16, r=0*.*866*)). On the other hand, the coefficients associated with *drs* null mutant were mostly positive for both bacteria types (Figure 5F *((Median: 0*.*596 (IQR: 0*.*359; 0*.*757); One-way Wilcoxon test for median> 0: <2*.*2e-16, r= 0*.*845*) and Figure 5H (*Median: 2*.*055 (IQR: 1*.*897; 2*.*192); One-way Wilcoxon test for median> 0: <2*.*2e-16, r=0*.*866*)). These results suggest that, throughout metamorphosis, a process that is dependent on the increase of AMP expression at pupariation is necessary for controlling the persistence of bacteria originating from gut purge imperfections.

When ecdysone signalling activity is decreased in the fat body and haemocytes (*Cg[ts]>EcRDN)*, a similar pattern can be observed: there is a higher proportion of individuals with larger loads of persisting bacteria (Figure 6A-B). The resampling strategy revealed that *Cg[ts]>EcRDN* samples have positive slope values for both *Lactobacillus* (Figure 6C, *Median: 2*.*746 (IQR: 2*.*562; 2*.*930); One-way Wilcoxon test for median>0: <2*.*2e-16, r=0*.*866)* and *Acetobacter* (Figure 6D, *Median: 1*.*981 (IQR: 1*.*795; 2*.*184); One-way Wilcoxon test for median>0: <2*.*2e-16, r=0*.*866)*, which is in accordance with increased bacterial loads during metamorphosis. Importantly, *Mex>EcRDN* individuals do not have increased bacterial loads during metamorphosis, suggesting that the local immune response does not have a measurable antibacterial effect relatively to the systemic response that is still operating in these conditions (Supplementary Figure 7).

**Figure 6.**
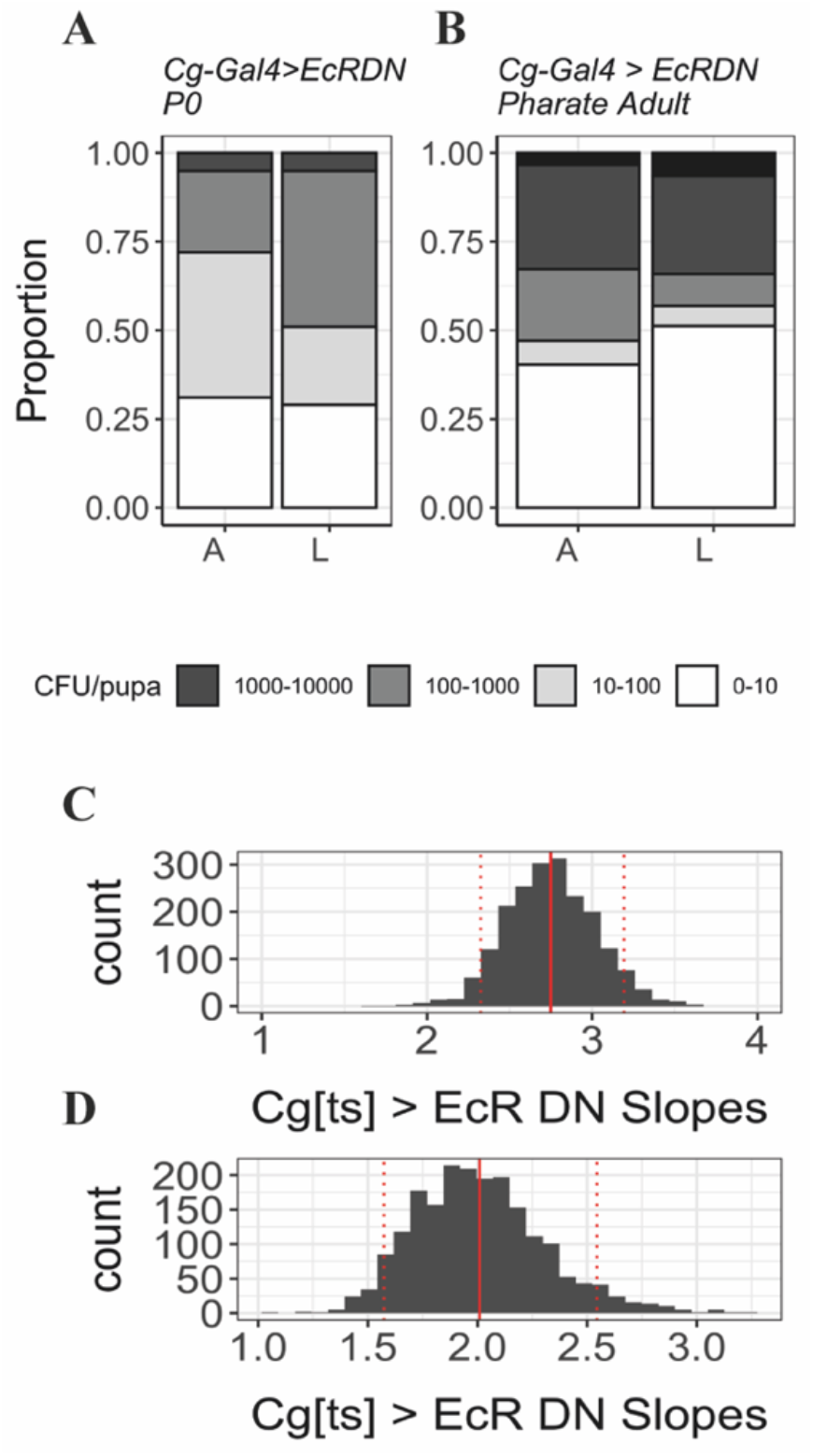
(A-D) Quantification in P0 pupae and pharate adults of *Lactobacillus* (L) and *Acetobacter* (A) with or without ecdysone signalling in the fat body and haemocytes. (A-B) When ecdysone sensitivity is decreased in fat body and haemocytes, the number of both bacteria types increases in P0 and pharate adults. Y-axis represents the proportion of individuals in each bacterial count category. (C-D) Coefficient (slopes) distribution in *Cg[ts]>EcRDN* samples of *Lactobacillus* (C) and *Acetobacter* (D). Doted lines represent the 0.05 and 0.95 quantiles; full line represents the mean slope.

## Discussion

In this study in *D. melanogaster*, we establish the association between ecdysone and the increased expression of *drs* and its paralogues at pupariation and the biological consequence of this process. We determine the functional link, to our knowledge yet undescribed in any species, between the endocrine and immune systems, acting both systemically and locally during metamorphosis to control the number of bacteria in the pupal case. The biological significance of the AMP peak at the onset of metamorphosis is revealed by the systemic expression of *drs*, as both mutant and ecdysone sensitivity-reduced individuals showed an increase rather than a decrease in the number of bacteria between P0 and pharate adult stages. Interestingly the effect was detected for both Gram-positive (*Lactobacillus*) and Gram-negative (*Acetobacter*) bacteria. Although Drs is generally characterized as an antifungal AMP, previous studies already demonstrated that it can also be induced by both Gram-positive and Gram-negative bacteria (35–41). As such, the unexpected higher impact of *drs* absence on *Acetobacter* may be explained by a direct impact on Gram-negative bacteria. Indeed, a previous study demonstrated that *drs* is induced in the fat body after a systemic infection with *Ecc15* (34), which suggests this AMP responds to the presence of Gram-negative bacteria and may be relevant under an infection scenario. Alternatively, the impact seen on Gram-negative bacteria might be due to some compensatory mechanism driven by other AMPs in the absence of Drs. Also of note that the humoral arm of defence is not the only one acting at this stage of development. Regan and colleagues demonstrated that at pupariation, ecdysone is required to activate haemocytes and to orchestrate the active migration of these cells towards wounding sites (6). Therefore, at this stage both ecdysone-regulated cellular and humoral mechanisms are likely to act in synergy to decrease the probability of a damaging infection.

Surprisingly two of the actors in this process, *drsl2* and *drsl5*, have not been found to have any antibacterial activity (34). However, both these AMPs are upregulated in the gut upon *Erwinia carotovora* infection, which suggests that they may be relevant during an immune challenge (34) and aligns with our findings. Further studies should ascertain the exact mode of action of these drosomycin-like peptides.

Because *drsl2* and *drsl5* are not proven to act as AMPs and given that the lack of local expression of *drsl2* did not affect bacterial persistence *per se*, we focused our efforts on disentangling more of the molecular regulation of *drs* expression in the fat body. We showed that the expression of *drs* at this stage depends upon the expression in the fat body of the ecdysone receptor target, *br-Z4*. This is in accordance to previous indications that Br is essential for the PGRP-LC-independent hormonal regulation of *drs* in S2 cells (4). However, these previous studies did not demonstrate this association *in vivo*, thus not establishing the spatial and temporal range of Br action on AMPs nor its relation to pupal viability. Furthermore, similar ecdysone-mediated immune activation through Br has been previously observed. Studies in the silkworm *Bombyx mori* showed that during metamorphosis, ecdysone activates the transcription of lysozyme via Br-Z2 (42) and of the AMP *lebocin* via Br-Z4 (32).

This intermediary role of Br in the regulation mechanism of ecdysone over the AMPs may explain why these effectors are only upregulated at pupariation, as revealed by our expression profile data. Although ecdysone controls all the moults throughout development, *drs* expression increases in response to this hormone specifically at pupariation, when Br is expressed in Holometabola (25,43). The regulation of AMPs by Br is also fecund regarding the interpretation of AMP expression throughout development between holo- and hemimetabolan insects. Johnston and colleagues demonstrated that these two insect groups have different AMP expression patterns, with holometabolans showing a strong upregulation of immunity that coincides with metamorphosis timing and Br expression(44). In contrast, hemimetabolans do not undergo a dramatic increase in immunity, possibly correlating with much less drastic changes in body remodelling during progressive metamorphosis. In accordance, there is indication that insects with complete metamorphosis suffer a reduction in density, diversity and altered composition of bacterial populations during metamorphosis (45–47), whereas those with incomplete metamorphosis show a constant increase in microbial density and diversity throughout development (48). These data, as ours, are consistent with the previously proposed hypothesis that complete metamorphosis elicits a prophylactic immune response (15,16). However, until now it was unclear how ecdysone could temporally regulate the expression of AMPs at pupariation.

Alterations in immunity at life stages with higher risk of infection seem to be a general feature of development, finding some parallels in both vertebrates and invertebrates. In *Drosophila triauraria*, genes of the *drs* family are induced during diapause, which is also considered to be a more susceptible period (49). Most interestingly, data from *Xenopus* also point towards an association between the timing of metamorphosis and a shift in the immune response (50). For example, Major Histocompatibility Complex I (MHC class I) molecules, which bind to peptide fragments derived from pathogens, are first detected at the beginning of metamorphosis (51) and the expression of MHC class II molecules is not triggered if metamorphosis is chemically blocked (52).

Together, our data suggest a regulatory mechanism through which ecdysone co-opted the immune system possibly to enhance the success of metamorphic processes. This study reveals a novel association between the endocrine and immune systems in *D. melanogaster*, which act both systemically and locally during metamorphosis. This recruitment of immunity mechanisms by the holometabolan metamorphosis programme presents other interesting features. Three AMPs explain around 95% of the peak observed at the L3 to pupal transition. Interestingly, these three AMPs have been co-opted differently and complementarily to produce, what we interpret as a full-fledged immune prophylactic response despite the absence of solid evidence of an antibacterial role for *drsl2* and *drsl5*. However, this speculation is tempting. *Drsl2*, was recruited to one of the main exposed surface epithelia of larvae, the midgut, that undergoes extensive remodelling at metamorphosis and *drs* is produced in the fat body at metamorphosis initiation, exerting its antibacterial action in a systemic fashion. These two mechanisms, with local and systemic action, share ecdysone control and account for close to 70% of the pupariation AMP peak.

This multi-layered developmentally-regulated immune response at the onset of pupariation underscores the sophistication and versatility of the co-option process (53–59) and hints at a possible contribution of this process to the success of metamorphosis as a whole.

## Materials and methods

### Drosophila stocks and husbandry

If not otherwise noticed, *w[1118]* flies were obtained from Dr. Luís Teixeira (Instituto Gulbenkian de Ciência, Portugal) and used as the reference wild-type strain. The following lines were obtained from the Bloomington Drosophila Stock Center: *w;UAS-mCD8GFP* (#5137), *Cg-Gal4* (#7011), *tub-Gal80[ts]*(#7017), *w; UAS-Cas9 P2*(#58986), *w;UAS-brZ4* (#51193) and *w;UAS-dsBroad* (#104648). Several stocks were generously shared by our colleagues, Drs Lynn Riddiford, University of Washington, USA (*w;UAS-EcRADN TP3 W650A*), Marc Dionne, Imperial College London, UK (*Hml-Gal4*), François Leulier, IGFL, France (*Mex-Gal4*), Kim Rewitz, University of Copenhagen, Denmark (*AkhR-Gal4*), Bruno Lemaitre, EPFL, Switzerland (*iso;iso;Drs*^*r(w+)*^ and the respective background control). The rescue lines *w;UAS-mCD8GFP; P{UAS-br*.*Z4}37-6* and *w;UAS-EcRADN TP3 W650A; P{UAS-br*.*Z4}37-6* were generated in house at Instituto Gulbenkian de Ciência (Portugal) by the Fly Transgenesis Service (Dr Gastón Guilgur). Standardly raised flies were kept on food with the following composition (g/mL): 4,5% molasse, 7,5% sugar, 7% cornmeal, 2% yeast extract, 1% agar, 2,5% Nipagin 10%. All fly lines we kept at 25°C in a 12L:12D photoperiod. For the ecdysone decreased sensitivity experiment and tissue specific CRISPR, animals were maintained at 17°C until the moult to the L3 and then transferred to 29°C to activate Gal4 activity.

### CRISPR knock-out flies

The tissue-specific UAS *br* CRISPR knockout constructs were cloned into pCFD6 vector (www.crisprflydesign.org, Addgene #73915) according to manufacturer’s instructions. Four independent gRNA constructs were cloned to target introns (see primers listed in S9 Table) of the *br* core region genomic DNA sequence for *br* common construct and Z4 isoform specific sequence for *br-Z4* construct to target only native genes but not transgenes. After sequenced, the constructs were inserted on either the second chromosome in a recipient line carrying phiC31 integrase and attP40 landing cite with an in-house modified line (*y,w, P{y+*.*nos-int*.*NLS}; P{CaryP y+}attP40*) or the third chromosome in a recipient line carrying phiC31 integrase and attP2 landing site, *y,w, P(y[+]*.*nos-int. NLS); P(CaryP)attP2* (gift from Dr. Diogo Manoel, Sidra Medical and Research Center, Qatar). Transgenesis was carried out in-house at Instituto Gulbenkian de Ciência (Portugal) by the Fly Transgenesis Service (Dr Gastón Guilgur).

### Larval and pupal staging

Larvae were resynchronized every 2h and kept in normal food for the appropriate time before collection in Trizol. Egg collections were performed on standard *Drosophila* food plates at 25°C, unless mentioned otherwise. All larvae used for this study were resynchronised either at the onset of the L2 or L3, as previously described (60). Eggs were collected every 4 hours and larval densities kept around 200 larvae/60 mm diameter plate. Newly moulted L2 or L3 larvae were collected every 2 hours. Collected larvae (up to a maximum of 30 larvae/vial) were raised in vials with standard food at 25°C for an additional twenty-four hours. Staged larvae were collected in Trizol for RNA extraction and stored at -80°C until use. For collection of white pre-pupae (P0), bottles with L3 larvae were monitored every 20 minutes, between 48 and 72h after the moult to the L3. Across this study, P0 were identified as motionless pre-pupae with evaginated anterior spiracles. After collection, samples were immediately stored in Trizol and stored at -80°C until RNA extraction.

### P0 and pharate adult collection and plating

P0 samples were collected as aforementioned. Pharate adults were identified as fully developed adults (with tanned wings and pigmented eyes) still to eclose. Pharate adults were dissected out of the pupal case and collected in 20 minutes intervals 48h to 72h after P0 collection. All samples were surface sterilized once with Milli-Q water, 70% ethanol and 13% bleach and rinsed (3x) with sterile water to eliminate bleach traces. After these treatments, each pupa was macerated in 300 μL of LB medium, of which 50 μL were spread onto MRS or D-Mannitol plates. The plates were incubated at 25°C for 48h for bacterial growth. For each stage and medium a sample size of 100 pupae/pharate was collected per genotype.

### Statistical analysis for bacterial growth across stages

To compare the pattern of variation between P0 and pharate adults across genotypes, we followed a resampling strategy (61): the initial sample was resampled in subsets of 50 entries, with no replacement, 2000 times, generating a simulated dataset. In each cycle, a negative binomial regression (*glm*.*nb(datat$cfu_pupae ∼ data$Stage)*) – which does not assume the variance and the mean of the data to be equivalent - was performed to model the change of bacteria according to the stage, generating a regression coefficient, or slope.

At the end of the 2000^th^ cycle, a set of these coefficients was obtained for each genotype. This strategy allows us to estimate a set of values that the negative binomial generalized linear regressions may assume, instead of getting just one value. In turn, this set can allow us to explore location and variation statistics, as well as better estimate the parameters of the sampling distribution of the coefficients, with a better notion of its precision.

Subsequently, we assessed if the central tendency of the population is greater or less than 0 using one-sided hypothesis tests. In parallel, a measure of the effect size was also calculated. Following a Shapiro-Wilk test of normality, a parametric or non-parametric approach for hypothesis testing and effect size calculation was applied. All data was analysed and all graphics were generated using Rstudio 1.2.5033.

### Germ-free flies

Newly laid eggs were collected into a sterile embryo basket and washed with autoclaved MilliQ water. The eggs were dechorionated and sterilised using a 13% Bleach solution (Sigma Aldrich®) for 10 minutes, washed again with Milli-Q water and sterilized using a 1% Virkon solution for 2 minutes. The sterilized embryos were transferred to axenic media with or without antibiotics (200 mg/mL Rifampicin, 100 mg/mL Tetracycline, 100 mg/mL Streptomycin, 15 mg/mL gentamicin). The axenic media contained all the normal ingredients of fly food, is autoclaved prior to use and supplemented with the fungicide Bavistin, which guarantees absence of any traces of microorganisms, including yeast. All plates and bottles were kept inside sterilised containers at 25°C and all manipulations were conducted in a horizontal flood hood, using sterile materials. After the treatment, the germ-free status was validated by plating surface-sterilized adult flies (from both germ free and control conditions) onto MRS and D-Mannitol media.

### RNA extraction, cDNA synthesis and real-quantitative PCR

RNA was isolated using the Direct-zol™ RNA MiniPrep Kit (Zymo®), following manufacturer’s instructions. DNase I treatment and cDNA synthesis were preformed using the RQ1 RNASE-FREE DNASE 1* (Promega®) and the RevertAid H Minus First Strand cDNA Synthesis Kit (Thermo Scientific®), respectively. qPCR was performed using the SYBR® Green PCR Master Mix (Applied Biosystems®) and ABI 7900HT (Applied Biosystems). The PCR conditions used in all experiments were: initial denaturation/ enzyme activation, 95°C for 10’; followed by 45 cycles of denaturation, 95°C for 10”; annealing, 60°C for 10”; extension, 72°C for 30”. Specificity of PCR amplification was verified by melting curve: 95°C for 10”, 65°C for 1’, 97°C for 1”; cooling, 37°C, 30”. Primers used for all qPCR reactions are represented in S10 Table. Foldchange was determined using the ΔΔCT method, which compares the expression of a given gene of interest to a housekeeping gene between experimental and control samples (62,63). Control L3 samples were used as reference. The data was analysed with a general linear model, using the genotype, stage and gene as explicable variables, as follows: glm(log(Foldchange)∼Genotype*Stage*Gene, data=mydata). The expression level for each gene at P0 and pharate was compared within and between genotypes through multiple comparisons using the *lsmeans* package from Rstudio 1.2.5033.

### Br binding sites prediction

The DNA binding motif matrix for Br-Z4 was obtained from the JASPAR database (http://jaspar.genereg.net) and the 5’ 2kb region of the *drosomycin* gene was acquired in Flybase (http://flybase.org). Both sequences were uploaded in FIMO software (http://meme-suite.org/tools/fimo) (33), using a p-value < 0.0001 as a filter.

## Acknowledgements

We thank Drs Lynn Riddiford, Marc Dionne, Luís Teixeira, Bruno Lemaitre, François Leulier, Kim Rewitz and Pedro Domingos for stock sharing. We thank Liliana Vieira and Sandra Crisóstomo for technical assistance, Gastón Guilgur for transgenesis services and André Barros for help with statistical analysis. Stocks obtained from the Bloomington Drosophila Stock Center (NIH P40OD018537) were used in this study. This work was supported by: Instituto Gulbenkian de Ciência/Fundação Calouste Gulbenkian; Fundação para a Ciência e a Tecnologia (FCT, Portugal) to CN (PD/BD/114344/2016); CONGENTO, project LISBOA-01-0145-FEDER-022170, co-financed by Lisboa Regional Operational Programme (Lisboa 2020), under the Portugal 2020 Partnership Agreement, through the European Regional Development Fund (ERDF), and Fundação para a Ciência e a Tecnologia (FCT, Portugal).

## Author Contributions

CN, Conceptualization, Formal analysis, Investigation, Visualization, Methodology, Writing—original draft, review and editing;

TK, Conceptualization, Investigation, Methodology, Writing—review and editing;

ES, Conceptualization, Supervision, Investigation, Funding acquisition, Methodology, Writing— original draft, review and editing.

## Supplementary Figures and Tables

### Supplementary Figures

**Supplementary Figure 1.**
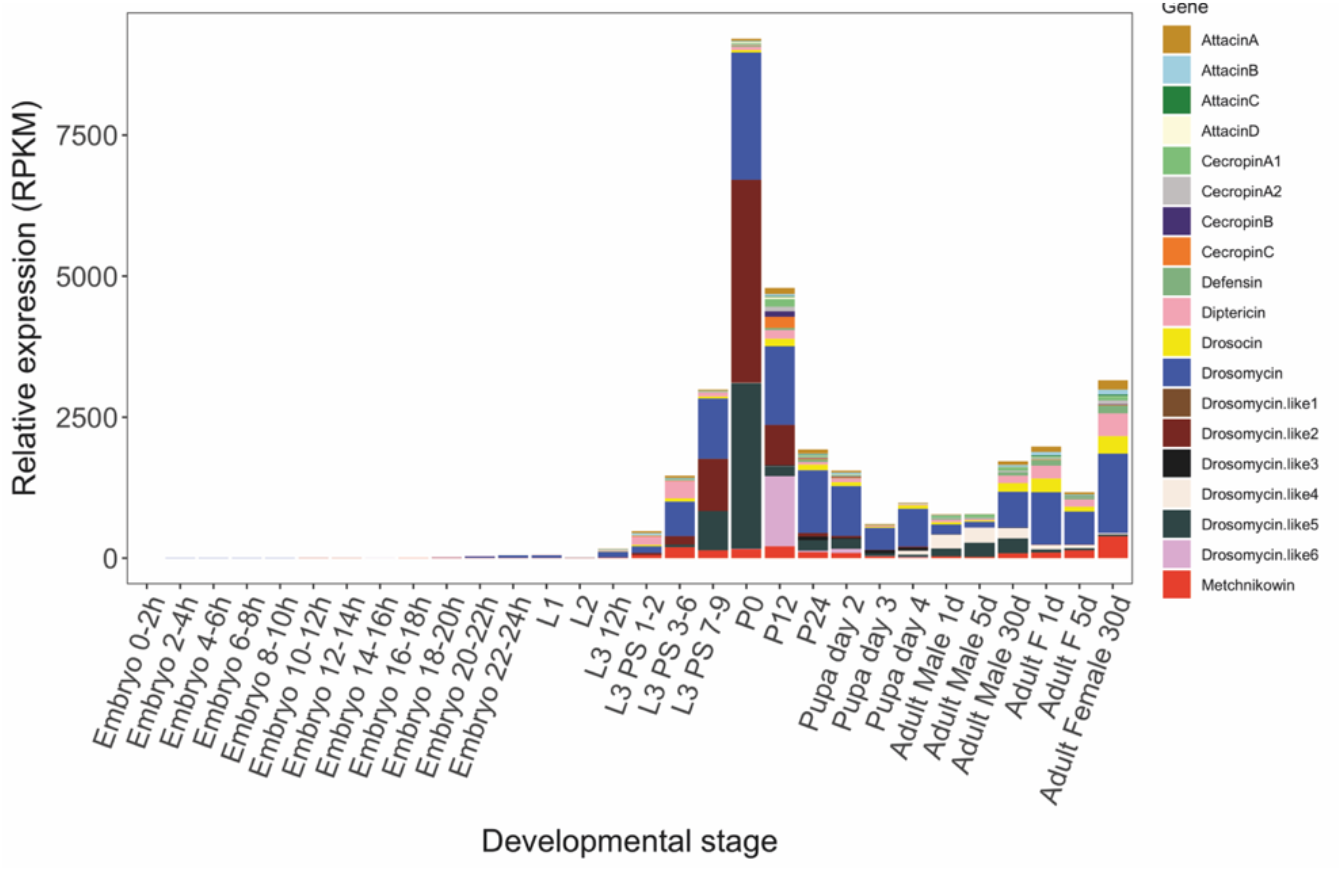
High-throughput temporal AMP expression. Expression refers to whole body, with levels represented as linear and scaled to maximum expression (publicly available on Flybase http://flybase.org/ (Thurmond et al., 2019)).

**Supplementary Figure 2.**
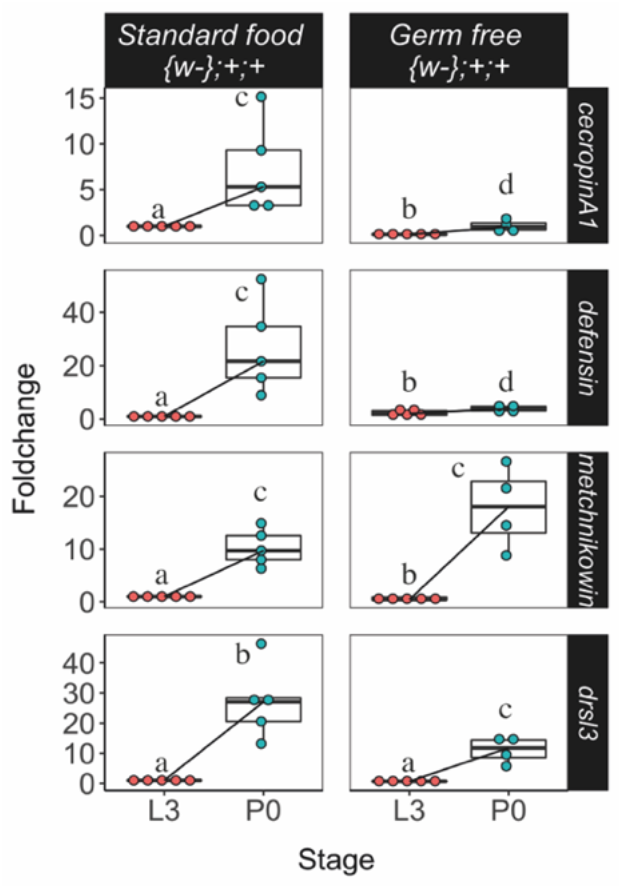
The expression levels of *cecropin A1, defensin, metchnikowin* and *drsl3* increase at pupariation and are not lost in GF conditions, albeit to different degrees.

**Supplementary Figure 3-1.**
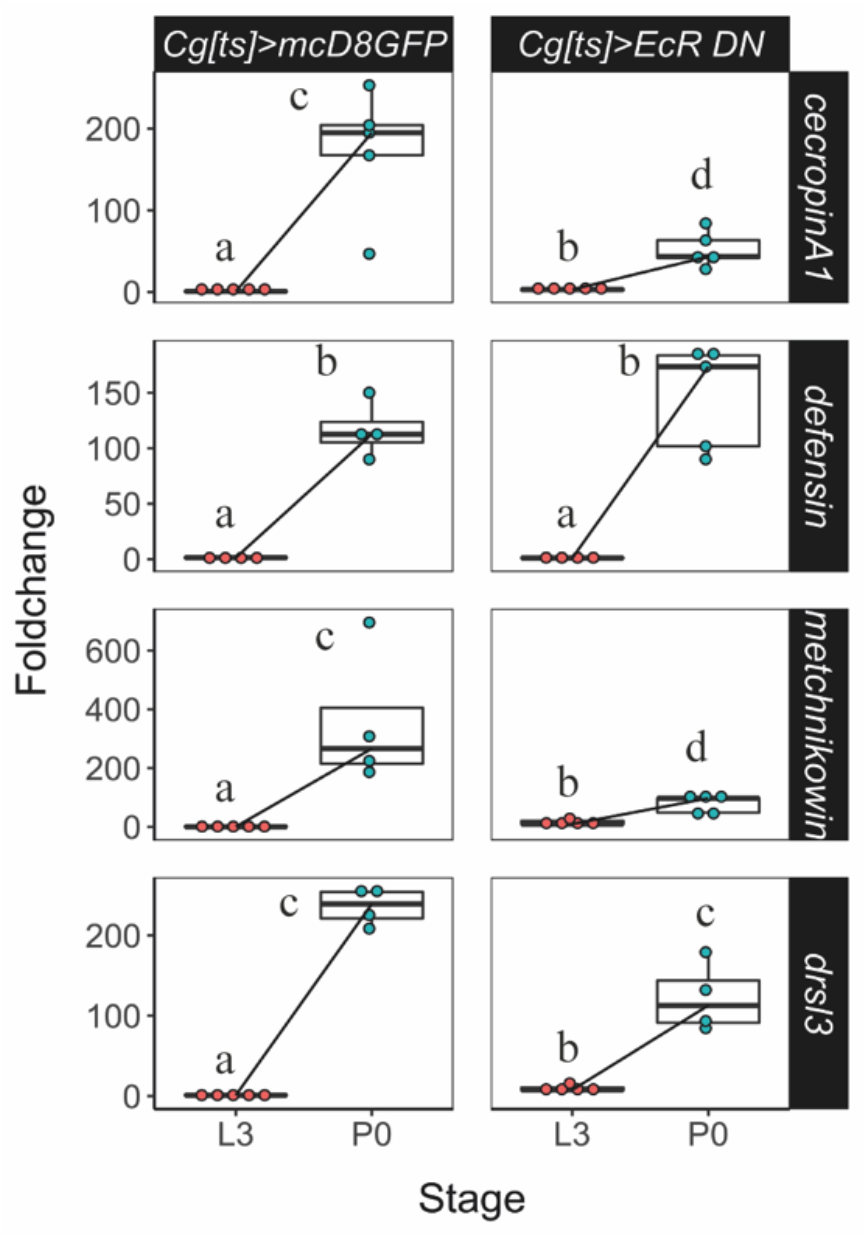
Reduced expression of ecdysone sensitivity in the fat body and haemocytes does not affect the expression of *cecropin A1, defensin, metchnikowin* and *drsl3*.

**Supplementary Figure 3-2.**
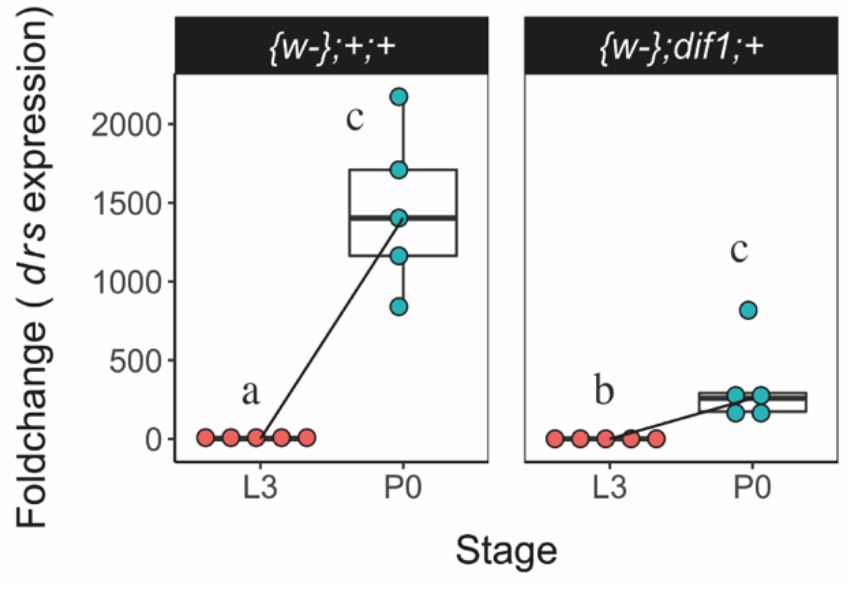
*drs* expression increases at P0 in *{w-};dif1;+* mutant.

**Supplementary Figure 3-3.**
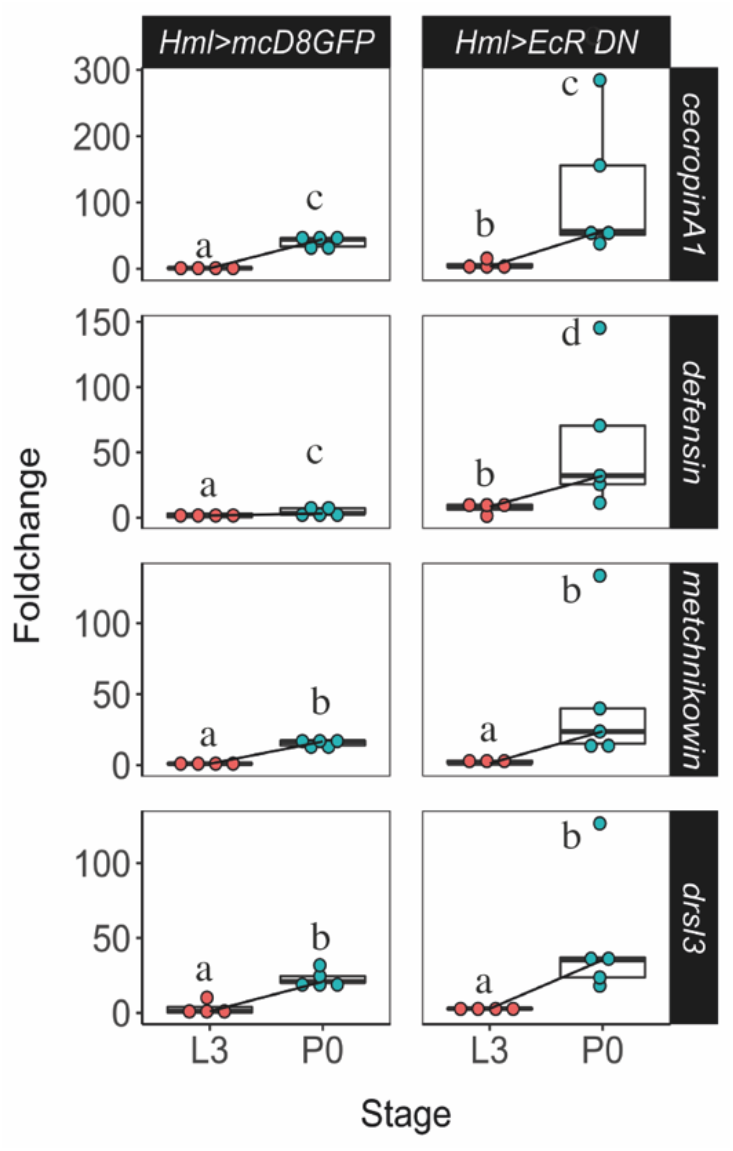
Reduced ecdysone sensitivity in the haemocytes does not affect the expression of of *cecropin A1, defensin, metchnikowin* and *drsl3*.

**Supplementary Figure 3-4.**
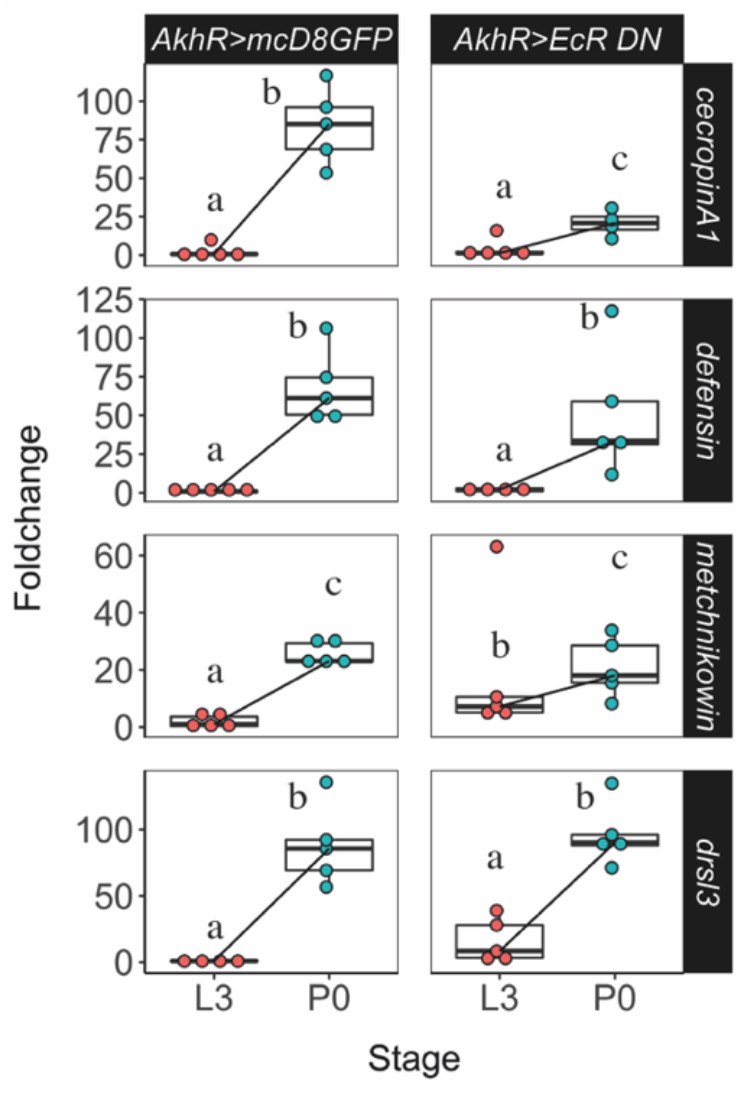
The expression of *cecropin A1, defensin, metchnikowin* and *drsl3* is unaffected by decreased ecdysone sensitivity specifically in the fat body.

**Supplementary Figure 4.**
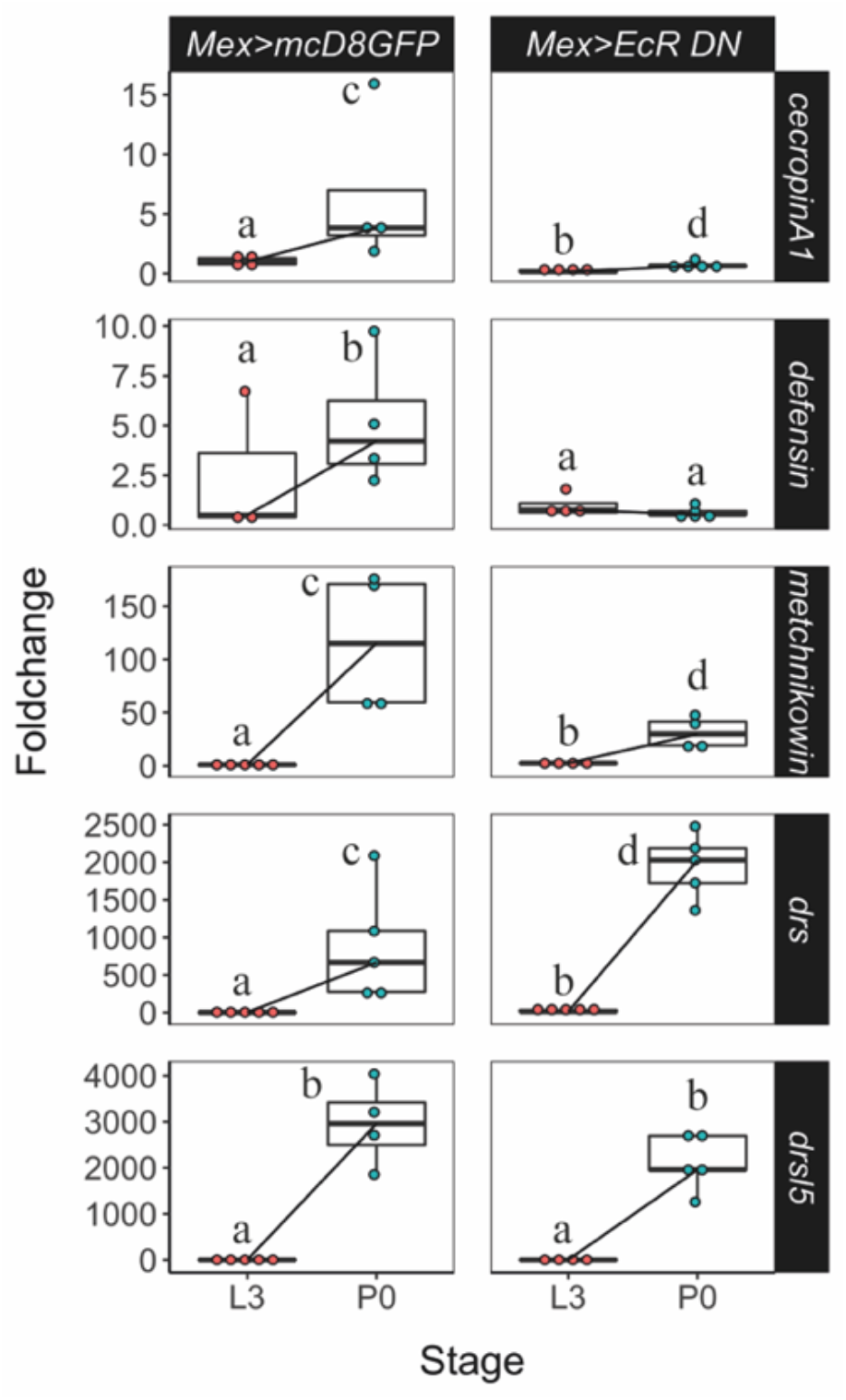
Decreased ecdysone signalling in the midgut results in significant decrease in mRNA levels for: *cecropin A1* (*lsmeans(pairwise∼Genotype*|*P0)<0*.*0001; lsmeans(pairwise∼Stage*|*Mex>EcRDN)= 0*.*0038), defensin (lsmeans(pairwise∼Genotype*|*P0) <0*.*0001; lsmeans(pairwise∼Stage*|*Mex>EcRDN)=0*.*3866*), *metchnikowin* (*lsmeans (pairwise∼Genotype*|*P0)=0*.*0060; lsmeans (pairwise∼Stage*|*Mex>EcRDN<0*.*0001*), and *drsl3 ((lsmeans(pairwise∼Genotype*|*P0)=0*.*0008; lsmeans(pairwise∼Stage*|*Mex>EcRDN)=0*.*2084*).

**Supplementary Figure 5-1.**
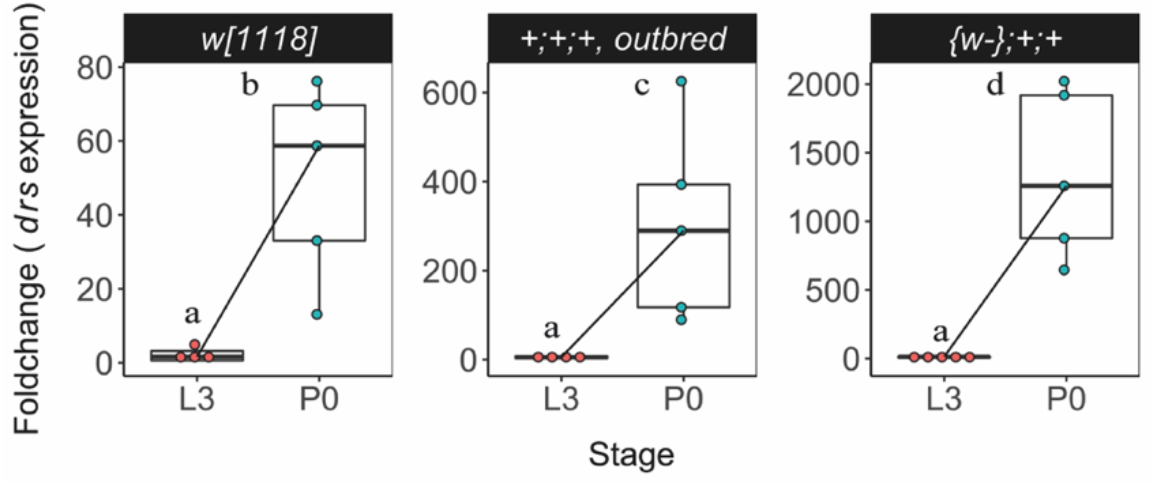
The expression of *drs* increased at pupariation regardless of the genetic background (*lsmeans(pairwise∼Stage*|*w1118)<0*.*0001; lsmeans(pairwise∼Stage*|*outbred <0*.*0001; lsmeans(pairwise∼Stage*|*Iso)<0*.*0001*). Each dot represents a sample of five pooled individuals; the lines connect the median of the samples at L3 and P0; different letters represent statistically significant differences in fold-change. Fold-change was determined using the ΔΔCT method.

**Supplementary Figure 5-2.**
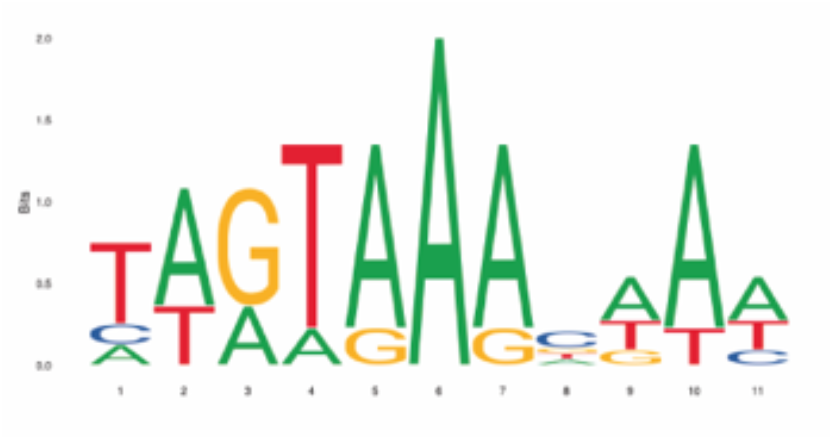
JASPAR DNA binding motif MA0013.1 for Br-Z4

**Supplementary Figure 5-3.**
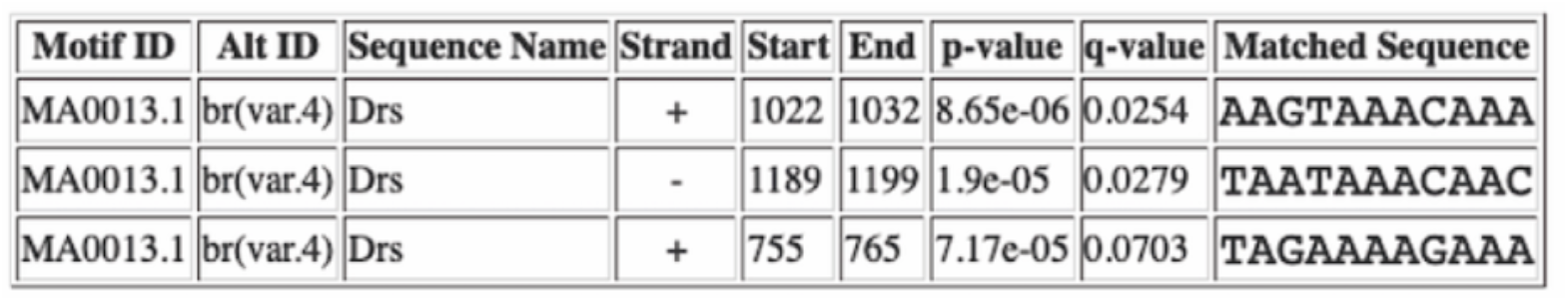
Br-Z4 binding motifs predicted by FIMO [29] software (http://meme-suite.org/tools/fimo) to be present in 2kb region of the 5’-end of the *drs* gene, using a p-value<0.0001 as filter.

**Supplementary Figure 6.**
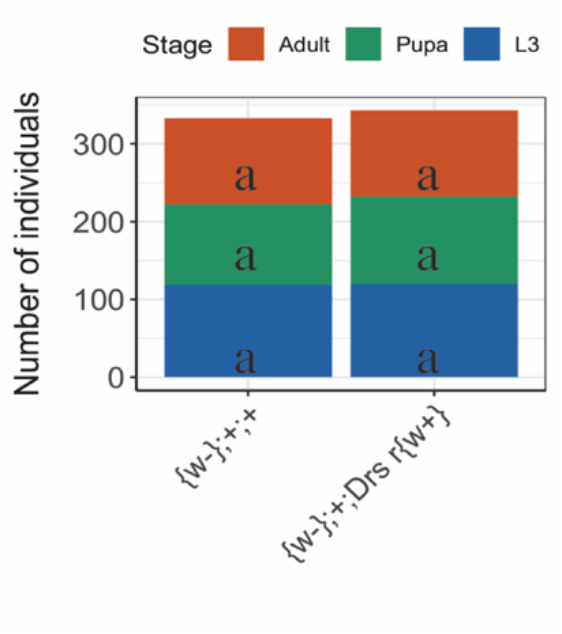
Control ({*w-};+;+*) and *drs* mutant ({*w-};+; Drs r{w+}*) lines showed no differences in mortality between L3, pupae and adults (*lsmeans(pairwise∼Stage*|*Genotype) <0*.*0001; lsmeans(pairwise∼Genotype*|*Stage)<0*.*0001*).

**Supplementary Figure 7.**
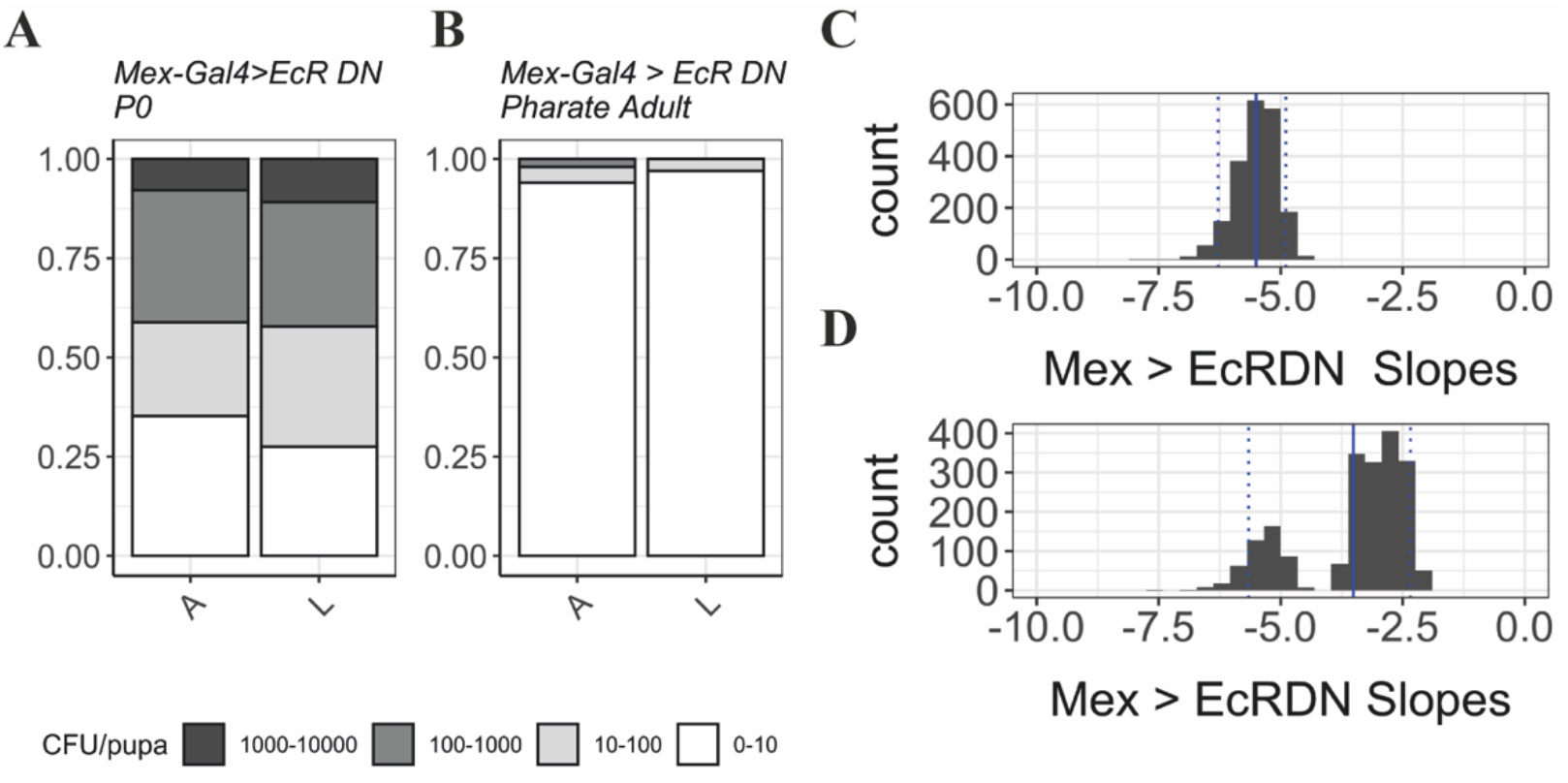
(A-B) Quantification in P0 pupae and pharate adults of *Lactobacillus* (L), growing in selective MRS medium, and *Acetobacter* (A), growing on selective D-Mannitol. The number of individuals with more bacteria is greater in P0 samples (A) than in pharate adults (B). Y-axis represents the proportion of individuals in each bacterial count category. (C-D) Coefficient (slopes) distribution in *Mex>EcRDN* samples of *Lactobacillus* (C) (*Median: -5*.*455 (IQR: -6*.*279; -4*.*895); One-way Wilcoxon test for median<0: <2*.*2e-16, r=0*.*866*) and *Acetobacter* (D) (*Median: -3*.*134 (IQR: -5*.*656; -2*.*342); One-way Wilcoxon test for median>0: <2*.*2e-16, r=0*.*866)*. Doted lines represent the 0.05 and 0.95 quantiles; full line represents the mean slope.

### Supplementary Tables

**S1 Table.**
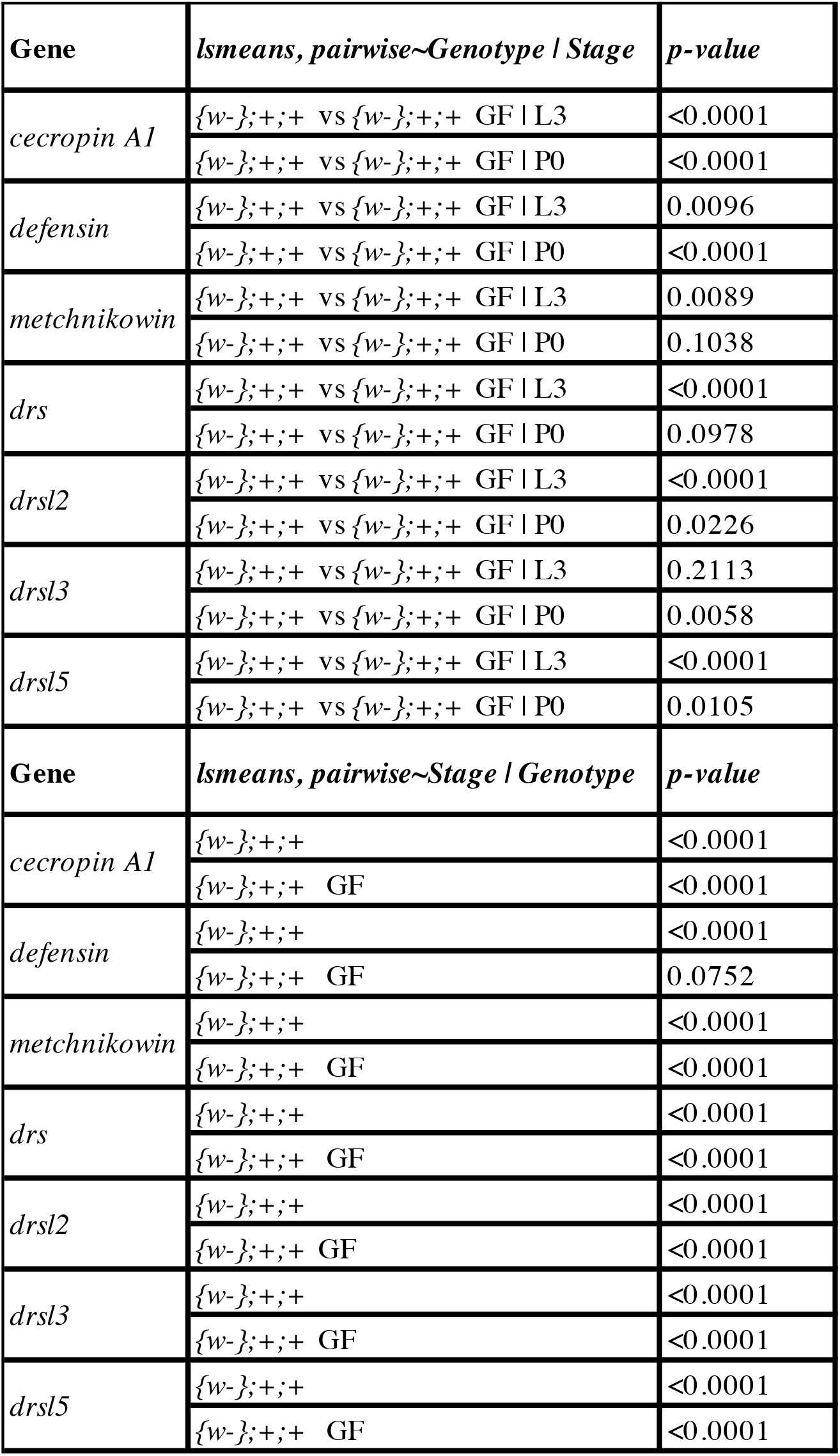
Statistical analysis of AMPs expression in an isogenic line (*{w-};+;+*) in normal and germ-free (GF) conditions at L3 and P0 stages. lsmeans, pairwise∼Genotype | Stage represents differences within the same stage, between genotypes. lsmeans, pairwise∼Stage | Genoype represents the difference between L3 and P0 within each genotype.

**S2 Table.** Statistical analysis of AMP expression in samples with decreased ecdysone sensitivity in the fat body and haemocytes (*Cg>EcRDN*). lsmeans, pairwise∼Genotype | Stage represents differences within the same stage, between genotypes. lsmeans, pairwise∼Stage | Genotype represents the difference between L3 and P0 within each genotype.

| Gene | lsmeans, pairwise~Genotype \| Stage | p-value |
| --- | --- | --- |
| *cecropin A1* | *Cg[ts]>mcD8GFP* vs *Cg[ts]>EcR DN* \| L3 | 0.0057 |
|  | *Cg[ts]>mcD8GFP* vs *Cg[ts]>EcR DN* \| P0 | 0.0064 |
| *defensin* | *Cg[ts]>mcD8GFP* vs *Cg[ts]>EcR DN* \| L3 | 0.8361 |
|  | *Cg[ts]>mcD8GFP* vs *Cg[ts]>EcR DN* \| P0 | 0.6428 |
| *metchnikowin* | *Cg[ts]>mcD8GFP* vs *Cg[ts]>EcR DN* \| L3 | <0.0001 |
|  | *Cg[ts]>mcD8GFP* vs *Cg[ts]>EcR DN* \| P0 | 0.0012 |
| *drs* | *Cg[ts]>mcD8GFP* vs *Cg[ts]>EcR DN* \| L3 | 0.7558 |
|  | *Cg[ts]>mcD8GFP* vs *Cg[ts]>EcR DN* \| P0 | <0.0001 |
| *drsl2* | *Cg[ts]>mcD8GFP* vs *Cg[ts]>EcR DN* \| L3 | 0.6216 |
|  | *Cg[ts]>mcD8GFP* vs *Cg[ts]>EcR DN* \| P0 | 0.0192 |
| *drsl3* | *Cg[ts]>mcD8GFP* vs *Cg[ts]>EcR DN* \| L3 | <0.0001 |
|  | *Cg[ts]>mcD8GFP* vs *Cg[ts]>EcR DN* \| P0 | 0.1326 |
| *drsl5* | *Cg[ts]>mcD8GFP* vs *Cg[ts]>EcR DN* \| L3 | <0.0001 |
|  | *Cg[ts]>mcD8GFP* vs *Cg[ts]>EcR DN* \| P0 | 0.4561 |
| Gene | lsmeans, pairwise~Stage \| Genotype | p-value |
| *cecropin A1* | *Cg[ts]>mcD8GFP* \| L3 vs P0 | <0.0001 |
|  | *Cg[ts]>EcR DN* \| L3 vs P0 | <0.0001 |
| *defensin* | *Cg[ts]>mcD8GFP* \| L3 vs P0 | <0.0001 |
|  | *Cg[ts]>EcR DN* \| L3 vs P0 | <0.0001 |
| *metchnikowin* | *Cg[ts]>mcD8GFP* \| L3 vs P0 | <0.0001 |
|  | *Cg[ts]>EcR DN* \| L3 vs P0 | <0.0001 |
| *drs* | *Cg[ts]>mcD8GFP* \| L3 vs P0 | <0.0001 |
|  | *Cg[ts]>EcR DN* \| L3 vs P0 | 0.0017 |
| *drsl2* | *Cg[ts]>mcD8GFP* \| L3 vs P0 | <0.0001 |
|  | *Cg[ts]>EcR DN* \| L3 vs P0 | <0.0001 |
| *drsl3* | *Cg[ts]>mcD8GFP* \| L3 vs P0 | <0.0001 |
|  | *Cg[ts]>EcR DN* \| L3 vs P0 | <0.0001 |
| *drsl5* | *Cg[ts]>mcD8GFP* \| L3 vs P0 | <0.0001 |
|  | *Cg[ts]>EcR DN* \| L3 vs P0 | <0.0001 |

**S3 Table.** Statistical analysis of AMP expression in samples with decreased ecdysone sensitivity in haemocytes (*Hml>EcRDN*). lsmeans, pairwise∼Genotype | Stage represents differences within the same stage, between genotypes. lsmeans, pairwise∼Stage | Genotype represents the difference between L3 and P0 within each genotype.

| Gene | lsmeans, pairwise~Genotype \| Stage | p-value |
| --- | --- | --- |
| *cecropin A1* | *Hml>mcD8GFP* vs *Hml>EcR DN* \| L3 | 0.0047 |
|  | *Hml>mcD8GFP* vs *Hml>EcR DN* \| P0 | 0.1124 |
| *defensin* | *Hml>mcD8GFP* vs *Hml>EcR DN* \| L3 | 0.0012 |
|  | *Hml>mcD8GFP* vs *Hml>EcR DN* \| P0 | <0.0001 |
| *metchnikowin* | *Hml>mcD8GFP* vs *Hml>EcR DN* \| L3 | 0.2988 |
|  | *Hml>mcD8GFP* vs *Hml>EcR DN* \| P0 | 0.1739 |
| *drs* | *Hml>mcD8GFP* vs *Hml>EcR DN* \| L3 | 0.0033 |
|  | *Hml>mcD8GFP* vs *Hml>EcR DN* \| P0 | 0.0185 |
| *drsl2* | *Hml>mcD8GFP* vs *Hml>EcR DN* \| L3 | 0.0512 |
|  | *Hml>mcD8GFP* vs *Hml>EcR DN* \| P0 | 0.8840 |
| *drsl3* | *Hml>mcD8GFP* vs *Hml>EcR DN* \| L3 | 0.0732 |
|  | *Hml>mcD8GFP* vs *Hml>EcR DN* \| P0 | 0.3020 |
| *drsl5* | *Hml>mcD8GFP* vs *Hml>EcR DN* \| L3 | 0.0065 |
|  | *Hml>mcD8GFP* vs *Hml>EcR DN* \| P0 | 0.1605 |
| Gene | lsmeans, pairwise~Stage \| Genotype | p-value |
| *cecropin A1* | *Hml>mcD8GFP* \| L3 vs P0 | <0.0001 |
|  | *Hml>EcR DN* \| L3 vs P0 | <0.0001 |
| *defensin* | *Hml>mcD8GFP* \| L3 vs P0 | 0.0118 |
|  | *Hml>EcR DN* \| L3 vs P0 | 0.0003 |
| *metchnikowin* | *Hml>mcD8GFP* \| L3 vs P0 | <0.0001 |
|  | *Hml>EcR DN* \| L3 vs P0 | <0.0001 |
| *drs* | *Hml>mcD8GFP* \| L3 vs P0 | <0.0001 |
|  | *Hml>EcR DN* \| L3 vs P0 | <0.0001 |
| *drsl2* | *Hml>mcD8GFP* \| L3 vs P0 | <0.0001 |
|  | *Hml>EcR DN* \| L3 vs P0 | <0.0001 |
| *drsl3* | *Hml>mcD8GFP* \| L3 vs P0 | <0.0001 |
|  | *Hml>EcR DN* \| L3 vs P0 | <0.0001 |
| *drsl5* | *Hml>mcD8GFP* \| L3 vs P0 | <0.0001 |
|  | *Hml>EcR DN* \| L3 vs P0 | <0.0001 |

**S4 Table.** Statistical analysis of AMP expression in samples with decreased ecdysone sensitivity in the fat body (*AkhR>EcRDN*). lsmeans, pairwise∼Genotype | Stage represents differences within the same stage, between genotypes. lsmeans, pairwise∼Stage | Genotype represents the difference between L3 and P0 within each genotype.

| Gene | lsmeans, pairwise~Genotype \| Stage | p-value |
| --- | --- | --- |
| *cecropin A1* | *AkhR>mcD8GFP* vs *AkhR>EcR DN* \| L3 | 0.2413 |
|  | *AkhR>mcD8GFP* vs *AkhR>EcR DN* \| P0 | 0.0228 |
| *defensin* | *AkhR>mcD8GFP* vs *AkhR>EcR DN* \| L3 | 0.4736 |
|  | *AkhR>mcD8GFP* vs *AkhR>EcR DN* \| P0 | 0.3747 |
| *metchnikowin* | *AkhR>mcD8GFP* vs *AkhR>EcR DN* \| L3 | 0.0001 |
|  | *AkhR>mcD8GFP* vs *AkhR>EcR DN* \| P0 | 0.5902 |
| *drs* | *AkhR>mcD8GFP* vs *AkhR>EcR DN* \| L3 | 0.0133 |
|  | *AkhR>mcD8GFP* vs *AkhR>EcR DN* \| P0 | <0.0001 |
| *drsl2* | *AkhR>mcD8GFP* vs *AkhR>EcR DN* \| L3 | 0.7280 |
|  | *AkhR>mcD8GFP* vs *AkhR>EcR DN* \| P0 | 0.4609 |
| *drsl3* | *AkhR>mcD8GFP* vs *AkhR>EcR DN* \| L3 | 0.0001 |
|  | *AkhR>mcD8GFP* vs *AkhR>EcR DN* \| P0 | 0.8518 |
| *drsl5* | *AkhR>mcD8GFP* vs *AkhR>EcR DN* \| L3 | 0.0058 |
|  | *AkhR>mcD8GFP* vs *AkhR>EcR DN* \| P0 | 0.3890 |
| Gene | lsmeans, pairwise~Stage \| Genotype | p-value |
| *cecropin A1* | *AkhR>mcD8GFP* \| L3 vs P0 | <0.0001 |
|  | *AkhR>EcR DN* \| L3 vs P0 | 0.0003 |
| *defensin* | *AkhR>mcD8GFP* \| L3 vs P0 | <0.0001 |
|  | *AkhR>EcR DN* \| L3 vs P0 | <0.0001 |
| *metchnikowin* | *AkhR>mcD8GFP* \| L3 vs P0 | <0.0001 |
|  | *AkhR>EcR DN* \| L3 vs P0 | 0.3266 |
| *drs* | *AkhR>mcD8GFP* \| L3 vs P0 | <0.0001 |
|  | *AkhR>EcR DN* \| L3 vs P0 | 0.6709 |
| *drsl2* | *AkhR>mcD8GFP* \| L3 vs P0 | <0.0001 |
|  | *AkhR>EcR DN* \| L3 vs P0 | <0.0001 |
| *drsl3* | *AkhR>mcD8GFP* \| L3 vs P0 | <0.0001 |
|  | *AkhR>EcR DN* \| L3 vs P0 | 0.0001 |
| *drsl5* | *AkhR>mcD8GFP* \| L3 vs P0 | <0.0001 |
|  | *AkhR>EcR DN* \| L3 vs P0 | <0.0001 |

**S5 Table.** Statistical analysis of AMP expression in samples with decreased ecdysone sensitivity in the migut (*Mex>EcRDN*). lsmeans, pairwise∼Genotype | Stage represents differences within the same stage, between genotypes. lsmeans, pairwise∼Stage | Genotype represents the difference between L3 and P0 within each genotype.

| Gene | Ismeans, pairwise~Genotype \| Stage | p-value |
| --- | --- | --- |
| *cecropin A1* | *Mex>mcD8GFP* vs *Mex>EcR DN* \| L3 | 0.0004 |
|  | *Mex>mcD8GFP* vs *Mex>EcR DN* \| P0 | <0.0001 |
| *defensin* | *Mex>mcD8GFP* vs *Mex>EcR DN* \| L3 | 0.7855 |
|  | *Mex>mcD8GFP* vs *Mex>EcR DN* \| P0 | <0.0001 |
| *metchnikowin* | *Mex>mcD8GFP* vs *Mex>EcR DN* \| L3 | 0.0371 |
|  | *Mex>mcD8GFP* vs *Mex>EcR DN* \| P0 | 0.0060 |
| *drs* | *Mex>mcD8GFP* vs *Mex>EcR DN* \| L3 | <0.0001 |
|  | *Mex>mcD8GFP* vs *Mex>EcR DN* \| P0 | 0.0076 |
| *drsl2* | *Mex>mcD8GFP* vs *Mex>EcR DN* \| L3 | 0.8069 |
|  | *Mex>mcD8GFP* vs *Mex>EcR DN* \| P0 | <0.0001 |
| *drsl3* | *Mex>mcD8GFP* vs *Mex>EcR DN* \| L3 | <0.0001 |
|  | *Mex>mcD8GFP* vs *Mex>EcR DN* \| P0 | 0.0008 |
| *drsl5* | *Mex>mcD8GFP* vs *Mex>EcR DN* \| L3 | 0.9832 |
|  | *Mex>mcD8GFP* vs *Mex>EcR DN* \| P0 | 0.4479 |
| Gene | Ismeans, pairwise~Stage \| Genotype | p-value |
| *cecropin A1* | *Mex>mcD8GFP* \| L3 vs P0 | 0.0010 |
|  | *Mex>EcR DN* \| L3 vs P0 | 0.0038 |
| *defensin* | *Mex>mcD8GFP* \| L3 vs P0 | 0.0028 |
|  | *Mex>EcR DN* \| L3 vs P0 | 0.3866 |
| *metchnikowin* | *Mex>mcD8GFP* \| L3 vs P0 | <0.0001 |
|  | *Mex>EcR DN* \| L3 vs P0 | <0.0001 |
| *drs* | *Mex>mcD8GFP* \| L3 vs P0 | <0.0001 |
|  | *Mex>EcR DN* \| L3 vs P0 | <0.0001 |
| *drsl2* | *Mex>mcD8GFP* \| L3 vs P0 | <0.0001 |
|  | *Mex>EcR DN* \| L3 vs P0 | <0.0001 |
| *drsl3* | *Mex>mcD8GFP* \| L3 vs P0 | <0.0001 |
|  | *Mex>EcR DN* \| L3 vs P0 | 0.2084 |
| *drsl5* | *Mex>mcD8GFP* \| L3 vs P0 | <0.0001 |
|  | *Mex>EcR DN* \| L3 vs P0 | <0.0001 |

**S6 Table.** Statistical analysis of AMP expression in samples with decreased ecdysone sensitivity in the tracheal system (*btl>EcRDN*). lsmeans, pairwise∼Genotype | Stage represents differences within the same stage, between genotypes. lsmeans, pairwise∼Stage | Genotype represents the difference between L3 and P0 within each genotype.

| Gene | lsmeans, pairwise~Genotype \| Stage | p-value |
| --- | --- | --- |
| *cecropin A1* | *Btl>mcD8GFP* vs *Btl>EcR DN* \| L3 | 0.0197 |
|  | *Btl>mcD8GFP* vs *Btl>EcR DN* \| P0 | 0.8417 |
| *defensin* | *Btl>mcD8GFP* vs *Btl>EcR DN* \| L3 | 0.1219 |
|  | *Btl>mcD8GFP* vs *Btl>EcR DN* \| P0 | 0.0336 |
| *metchnikowin* | *Btl>mcD8GFP* vs *Btl>EcR DN* \| L3 | <0.0001 |
|  | *Btl>mcD8GFP* vs *Btl>EcR DN* \| P0 | 0.6789 |
| *drs* | *Btl>mcD8GFP* vs *Btl>EcR DN* \| L3 | 0.0098 |
|  | *Btl>mcD8GFP* vs *Btl>EcR DN* \| P0 | 0.1636 |
| *drsl2* | *Btl>mcD8GFP* vs *Btl>EcR DN* \| L3 | 0.0197 |
|  | *Btl>mcD8GFP* vs *Btl>EcR DN* \| P0 | 0.9116 |
| *drsl3* | *Btl>mcD8GFP* vs *Btl>EcR DN* \| L3 | <0.0001 |
|  | *Btl>mcD8GFP* vs *Btl>EcR DN* \| P0 | 0.4955 |
| *drsl5* | *Btl>mcD8GFP* vs *Btl>EcR DN* \| L3 | 0.4510 |
|  | *Btl>mcD8GFP* vs *Btl>EcR DN* \| P0 | 0.1192 |
| Gene | lsmeans, pairwise~Stage \| Genotype | p-value |
| *cecropin A1* | *Btl>mcD8GFP* \| L3 vs P0 | <0.0001 |
|  | *Btl>EcR DN* \| L3 vs P0 | 0.0001 |
| *defensin* | *Btl>mcD8GFP* \| L3 vs P0 | 0.0027 |
|  | *Btl>EcR DN* \| L3 vs P0 | 0.0010 |
| *metchnikowin* | *Btl>mcD8GFP* \| L3 vs P0 | <0.0001 |
|  | *Btl>EcR DN* \| L3 vs P0 | 0.3711 |
| *drs* | *Btl>mcD8GFP* \| L3 vs P0 | <0.0001 |
|  | *Btl>EcR DN* \| L3 vs P0 | <0.0001 |
| *drsl2* | *Btl>mcD8GFP* \| L3 vs P0 | <0.0001 |
|  | *Btl>EcR DN* \| L3 vs P0 | <0.0001 |
| *drsl3* | *Btl>mcD8GFP* \| L3 vs P0 | <0.0001 |
|  | *Btl>EcR DN* \| L3 vs P0 | 0.0004 |
| *drsl5* | *Btl>mcD8GFP* \| L3 vs P0 | <0.0001 |
|  | *Btl>EcR DN* \| L3 vs P0 | <0.0001 |

**S7 Table.**
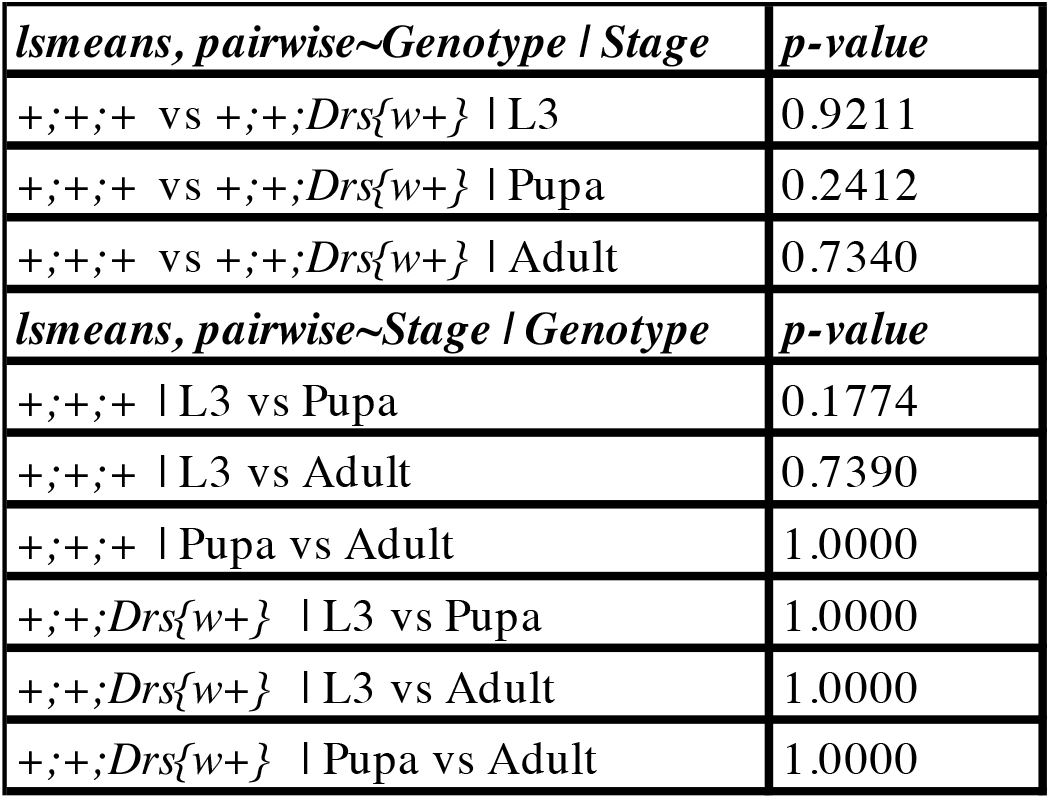
Statistical analysis of mortality between genotypes, across stages. lsmeans, pairwise∼Genotype | Stage represents differences within the same stage, between genotypes. lsmeans, pairwise∼Stage | Genotype represents the difference between L3 and P0 within each genotype.

**S8 Table.** Statistical analysis of AMP expression when *br* or *br-Z4* are knocked-out specifically in the fat body and haemocytes. lsmeans, pairwise∼Genotype | Stage represents differences within the same stage, between genotypes. L3 is coloured in red and P0 in blue. lsmeans, pairwise∼Stage | Genotype represents the difference between L3 and P0 within each genotype.

| **Gene** | **lsmeans, pairwise~Genotype \| Stage** | **p-value** |
| --- | --- | --- |
| *cecropin A1* | *Cg;Cas9>+,+,+ vs Cg;Cas9>br KO \| L3* | 1.0000 |
|  | *Cg;Cas9>+,+,+ vs Cg;Cas9>br-Z4 KO \| L3* | 1.0000 |
|  | *Cg;Cas9>br KO Cg;Cas9>br-Z4 KO \| L3* | 1.0000 |
|  | *Cg;Cas9>+,+,+ vs Cg;Cas9>br KO \| P0* | 1.0000 |
|  | *Cg;Cas9>+,+,+ vs Cg;Cas9>br-Z4 KO \| P0* | 1.0000 |
|  | *Cg;Cas9>br KO Cg;Cas9>br-Z4 KO \| P0* | 1.0000 |
| *defensin* | *Cg;Cas9>+,+,+ vs Cg;Cas9>br KO \| L3* | 0.6537 |
|  | *Cg;Cas9>+,+,+ vs Cg;Cas9>br-Z4 KO \| L3* | 1.0000 |
|  | *Cg;Cas9>br KO Cg;Cas9>br-Z4 KO \| L3* | 0.5756 |
|  | *Cg;Cas9>+,+,+ vs Cg;Cas9>br KO \| P0* | 0.2314 |
|  | *Cg;Cas9>+,+,+ vs Cg;Cas9>br-Z4 KO \| P0* | 0.0003 |
|  | *Cg;Cas9>br KO Cg;Cas9>br-Z4 KO \| P0* | 0.0831 |
| *metchnikowin* | *Cg;Cas9>+,+,+ vs Cg;Cas9>br KO \| L3* | 1.0000 |
|  | *Cg;Cas9>+,+,+ vs Cg;Cas9>br-Z4 KO \| L3* | 1.0000 |
|  | *Cg;Cas9>br KO Cg;Cas9>br-Z4 KO \| L3* | 0.7209 |
|  | *Cg;Cas9>+,+,+ vs Cg;Cas9>br KO \| P0* | 0.8651 |
|  | *Cg;Cas9>+,+,+ vs Cg;Cas9>br-Z4 KO \| P0* | 1.0000 |
|  | *Cg;Cas9>br KO Cg;Cas9>br-Z4 KO \| P0* | 0.4444 |
| *drs* | *Cg;Cas9>+;+;+ vs Cg;Cas9>br KO \| L3* | 1.0000 |
|  | *Cg;Cas9>+;+;+ vs Cg;Cas9>br-Z4 KO \| L3* | 1.0000 |
|  | *Cg;Cas9>br KO Cg;Cas9>br-Z4 KO \| L3* | 1.0000 |
|  | *Cg;Cas9>+;+;+ vs Cg;Cas9>br KO \| P0* | <0.0001 |
|  | *Cg;Cas9>+;+;+ vs Cg;Cas9>br-Z4 KO \| P0* | <0.0001 |
|  | *Cg;Cas9>br KO Cg;Cas9>br-Z4 KO \| P0* | 0.0002 |
| *drsl2* | *Cg;Cas9>+;+;+ vs Cg;Cas9>br KO \| L3* | 1.0000 |
|  | *Cg;Cas9>+;+;+ vs Cg;Cas9>br-Z4 KO \| L3* | 1.0000 |
|  | *Cg;Cas9>br KO Cg;Cas9>br-Z4 KO \| L3* | 1.0000 |
|  | *Cg;Cas9>+;+;+ vs Cg;Cas9>br KO \| P0* | 1.0000 |
|  | *Cg;Cas9>+;+;+ vs Cg;Cas9>br-Z4 KO \| P0* | 1.0000 |
|  | *Cg;Cas9>br KO Cg;Cas9>br-Z4 KO \| P0* | 1.0000 |
| *drsl3* | *Cg;Cas9>+;+;+ vs Cg;Cas9>br KO \| L3* | 1.0000 |
|  | *Cg;Cas9>+;+;+ vs Cg;Cas9>br-Z4 KO \| L3* | 1.0000 |
|  | *Cg;Cas9>br KO Cg;Cas9>br-Z4 KO \| L3* | 1.0000 |
|  | *Cg;Cas9>+;+;+ vs Cg;Cas9>br KO \| P0* | 1.0000 |
|  | *Cg;Cas9>+;+;+ vs Cg;Cas9>br-Z4 KO \| P0* | 0.9050 |
|  | *Cg;Cas9>br KO Cg;Cas9>br-Z4 KO \| P0* | 0.9168 |
| *drsl5* | *Cg;Cas9>+;+;+ vs Cg;Cas9>br KO \| L3* | 0.9520 |
|  | *Cg;Cas9>+;+;+ vs Cg;Cas9>br-Z4 KO \| L3* | 0.0167 |
|  | *Cg;Cas9>br KO Cg;Cas9>br-Z4 KO \| L3* | 0.2020 |
|  | *Cg;Cas9>+;+;+ vs Cg;Cas9>br KO \| P0* | 0.1613 |
|  | *Cg;Cas9>+;+;+ vs Cg;Cas9>br-Z4 KO \| P0* | 0.3062 |
|  | *Cg;Cas9>+;+;+ Cg;Cas9>br-Z4 KO \| P0* | 1.0000 |
| **Gene** | **lsmeans, pairwise~Stage \| Genotype** | **p-value** |
| *cecropin A1* | *Cg;Cas9>+;+;+ \| L3 vs P0* | <0.0001 |
|  | *Cg;Cas9>br KO \| L3 vs P0* | <0.0001 |
|  | *Cg;Cas9>br-Z4 KO \| L3 vs P0* | <0.0001 |
| *defensin* | *Cg;Cas9>+;+;+ \| L3 vs P0* | <0.0001 |
|  | *Cg;Cas9>br KO \| L3 vs P0* | <0.0001 |
|  | *Cg;Cas9>br-Z4 KO \| L3 vs P0* | <0.0001 |
| *metchnikowin* | *Cg;Cas9>+;+;+ \| L3 vs P0* | <0.0001 |
|  | *Cg;Cas9>br KO \| L3 vs P0* | <0.0001 |
|  | *Cg;Cas9>br-Z4 KO \| L3 vs P0* | <0.0001 |
| *drs* | *Cg;Cas9>+;+;+ \| L3 vs P0* | <0.0001 |
|  | *Cg;Cas9>br KO \| L3 vs P0* | 0.0001 |
|  | *Cg;Cas9>br-Z4 KO \| L3 vs P0* | 0.4691 |
| *drsl2* | *Cg;Cas9>+;+;+ \| L3 vs P0* | <0.0001 |
|  | *Cg;Cas9>br KO \| L3 vs P0* | <0.0001 |
|  | *Cg;Cas9>br-Z4 KO* \| L3 vs P0 | <0.0001 |
| *drsl3* | *Cg;Cas9>+;+;+* \| L3 vs P0 | <0.0001 |
|  | *Cg;Cas9>br KO* \| L3 vs P0 | <0.0001 |
|  | *Cg;Cas9>br-Z4 KO* \| L3 vs P0 | <0.0001 |
| *drsl5* | *Cg;Cas9>+;+;+* \| L3 vs P0 | <0.0001 |
|  | *Cg;Cas9>br KO* \| L3 vs P0 | <0.0001 |
|  | *Cg;Cas9>br-Z4 KO* \| L3 vs P0 | <0.0001 |

**S9 Table.**
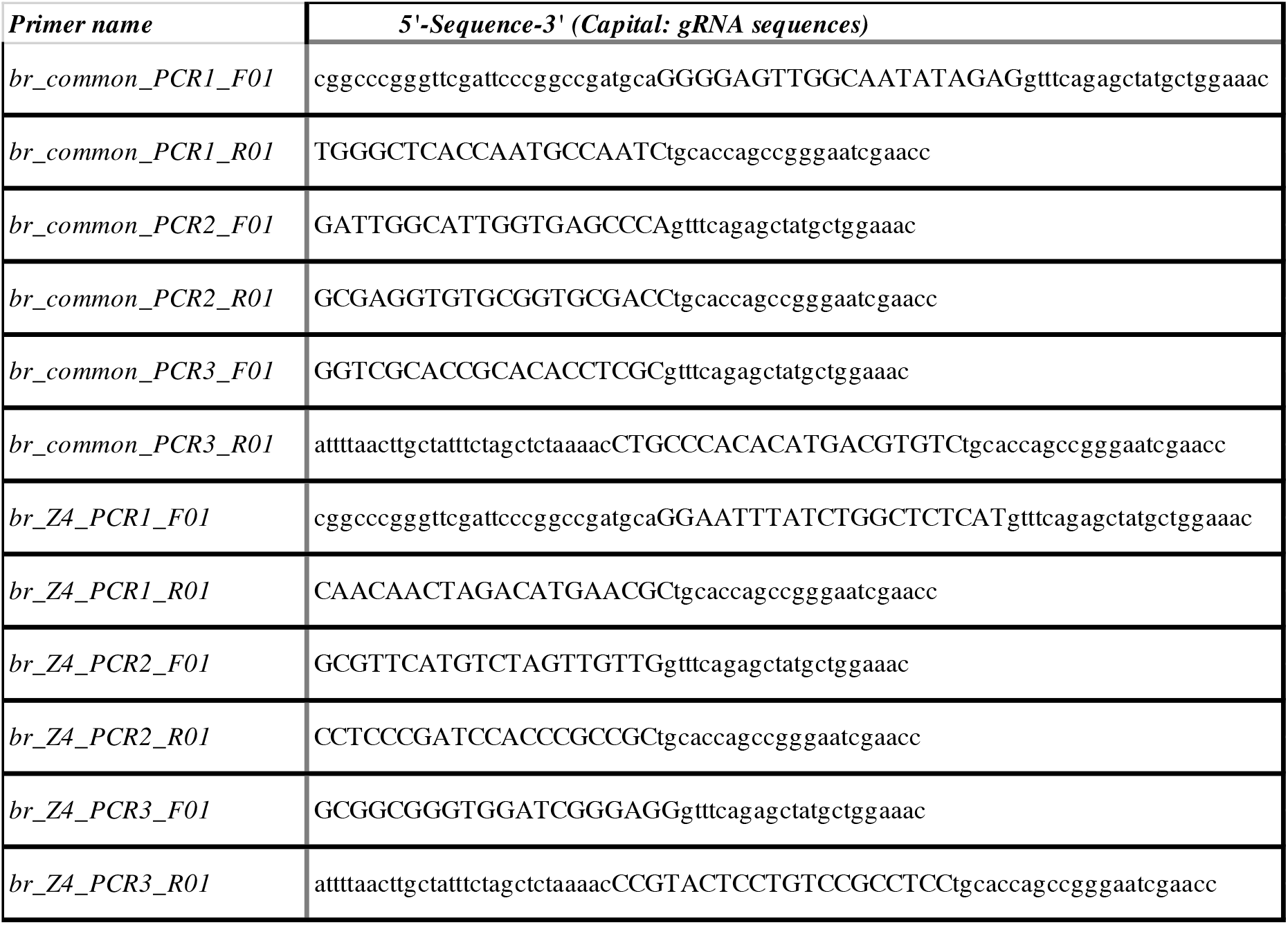
5’-3’ sequences of primers used for tissue specific CRISPR. Guide RNA (gRNA) sequences are represented in capital letters.

**S10 Table.**
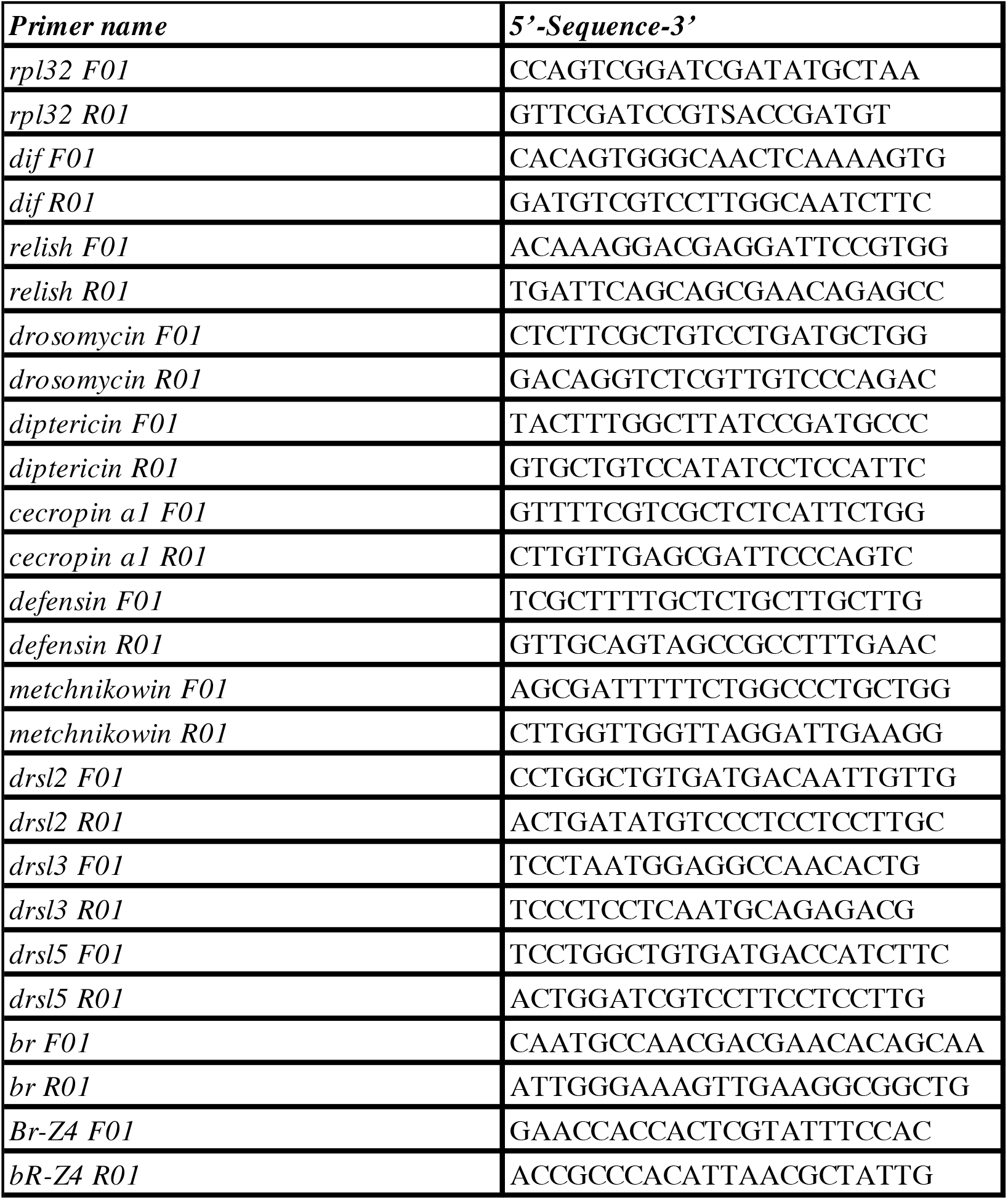
5’-3’ sequences of primers used for qPCR.

